# What is an archaeon and are the Archaea really unique?

**DOI:** 10.1101/256263

**Authors:** Ajith Harish

## Abstract

The recognition of the group Archaea as a major branch of the Tree of Life (ToL) prompted a new view of the evolution of biodiversity. The genomic representation of archaeal biodiversity has since significantly increased. In addition, advances in phylogenetic modeling of multi-locus datasets have resolved many recalcitrant branches of the ToL. Despite the technical advances and an expanded taxonomic representation, two important aspects of the origins and evolution of the Archaea remain controversial, even as we celebrate the 40th anniversary of the monumental discovery. These issues concern (i) the uniqueness (monophyly) of the Archaea, and (ii) the evolutionary relationships of the Archaea to the Bacteria and the Eukarya; both of these are relevant to the deep structure of the ToL. Here, to explore the causes for this persistent ambiguity, I examine multiple datasets that support contradicting conclusions. Results indicate that the uncertainty is primarily due to a scarcity of information in standard datasets — the core genes datasets — to reliably resolve the conflicts. These conflicts can be resolved efficiently by comparing patterns of variation in the distribution of functional genomic signatures, which are less diffused unlike patterns of primary sequence variation. Relatively lower heterogeneity in distribution patterns minimizes uncertainties, which supports statistically robust phylogenetic inferences, especially of the earliest divergences of life. This case study further highlights the limits of primary sequence data in resolving difficult phylogenetic problems and casts doubt on evolutionary inferences drawn solely from the analyses of a small set of core genes.

## Introduction

The recognition of the Archaea as the so-called “third form of life” was made possible in part by a new technology for sequence analysis, oligonucleotide cataloging, developed by Fredrik Sanger and colleagues in the 1960s (1, 2). Carl Woese’s insight of using this method, and the choice of the small subunit ribosomal RNA (16S/SSU rRNA) as a phylogenetic marker, not only put microorganisms on a phylogenetic map (or tree), but also revolutionized the field of molecular systematics that zukerkandl and Pauling had previously alluded to (3). Comparative analysis of organism-specific oligonucleotide signatures in SSU rRNA led to the recognition of a distinct group of microorganisms (2). Initially referred to as Archaeabacteria, these unusual organisms had ‘sequence signatures’ distinct from other bacteria (Eubacteria), and they were later found to be different from those of Eukarya (eukaryotes) as well. Many other features, including molecular, biochemical as well as ecological, corroborated the uniqueness of the Archaea. Thus the archaeal concept was established (2).

The study of microbial diversity and evolution has come a long way since then: sequencing microbial genomes, and directly from the environment without the need for culturing is now routine (4, 5). This wealth of sequence information is exciting not only for cataloging and organizing biodiversity, but also to understand the ecology and evolution of microorganisms – archaea and bacteria as well as eukaryotes – that make up a vast majority of the planetary biodiversity. Since large-scale exploration by the means of environmental genome sequencing became possible almost a decade ago, there has also been a palpable excitement and anticipation of the discovery of a fourth form of life or a “fourth domain” of life (6). The reference here is to a fourth form of cellular life, but not to viruses, which some have already proposed to be the fourth domain of the Tree of Life (ToL) (7). If a fourth form of life were to be found, what would the distinguishing features be, and how could it be measured, defined and classified?

Rather than the discovery of a fourth domain, and contrary to the expectations, however, current discussion is centered around the return to a dichotomous classification of life (8-10), despite the rapid expansion of sequenced biodiversity – hundreds of novel phyla descriptions (11, 12). The proposed dichotomous classification schemes, unfortunately, are in sharp contrast to each other, depending on: (i) whether the Archaea constitute a monophyletic group—a unique line of descent that is distinct from those of the Bacteria as well as the Eukarya; and (ii) whether the Archaea form a sister clade to the Eukarya or to the Bacteria. Both the issues stem from difficulties involved in resolving the deep branches of the ToL (9, 10, 13).

The twin issues, first recognized in the 80s based on single-gene (SSU rRNA) analyses (14), continue to be the subjects of a long-standing debate, which remains unresolved despite large-scale analyses of multi-gene datasets (4, 15-19). In addition to the choice of genes to be analyzed, the choice of the underlying character evolution model is at the core of contradictory results that either supports the Three-domains tree (4, 19) or the Eocyte tree (16, 18). In many cases, adding more data, either as enhanced taxon (species) sampling or enhanced character (gene) sampling, or both, can resolve ambiguities (20, 21). However, as the taxonomic diversity and evolutionary distance increases among the taxa studied, the number of conserved marker-genes that can be used for phylogenomic analyses decreases. Accordingly, resolving the phylogenetic relationships of the Archaea, Bacteria and Eukarya is restricted to a small set of genes—50 at most (18)—in spite of the large increase in the numbers of genomes sequenced and the associated development of sophisticated phylogenomic methods.

Based on a closer scrutiny of the phylogenomic datasets employed in the ongoing debate, I will show here that one of the reasons for this persistent ambiguity is that the ‘information’ necessary to resolve these conflicts is practically nonexistent in the standard marker-genes (i.e. core-genes) datasets employed routinely for phylogenomics. Further, I discuss analytical approaches that maximize the use of the information that is in genome sequence data and simultaneously minimize phylogenetic uncertainties. In addition, I discuss simple but important, yet undervalued, aspects of phylogenetic hypothesis testing, which together with the new approaches hold promise to resolve these long-standing issues effectively.

## Results

### Information in core genes is inadequate to resolve the archaeal radiation

Data-display networks (DDNs) are useful to examine and visualize character conflicts in phylogenetic datasets, especially in the absence of prior knowledge about the source of such conflicts (22, 23). While congruent data will be displayed as a tree in a DDN, incongruences are displayed as reticulations in the tree. Fig. 1A shows a neighbor-net analysis of the SSU rRNA alignment used to resolve the phylogenetic position of the recently discovered Asgard archaea (18). The DDN is based on character distances calculated as the observed genetic distance (p-distance) of 1,462 characters, and shows the total amount of conflict in the dataset that is incongruent with character bipartitions (splits). The edge (branch) lengths in the DDN correspond to the support for the respective splits. Accordingly, two well-supported sets of splits for the Bacteria and the Eukarya are observed. The Archaea, however, does not form a distinct, well-resolved/well-supported group, and is unlikely to correspond to a monophyletic group in a phylogenetic tree.

**Figure 1.**
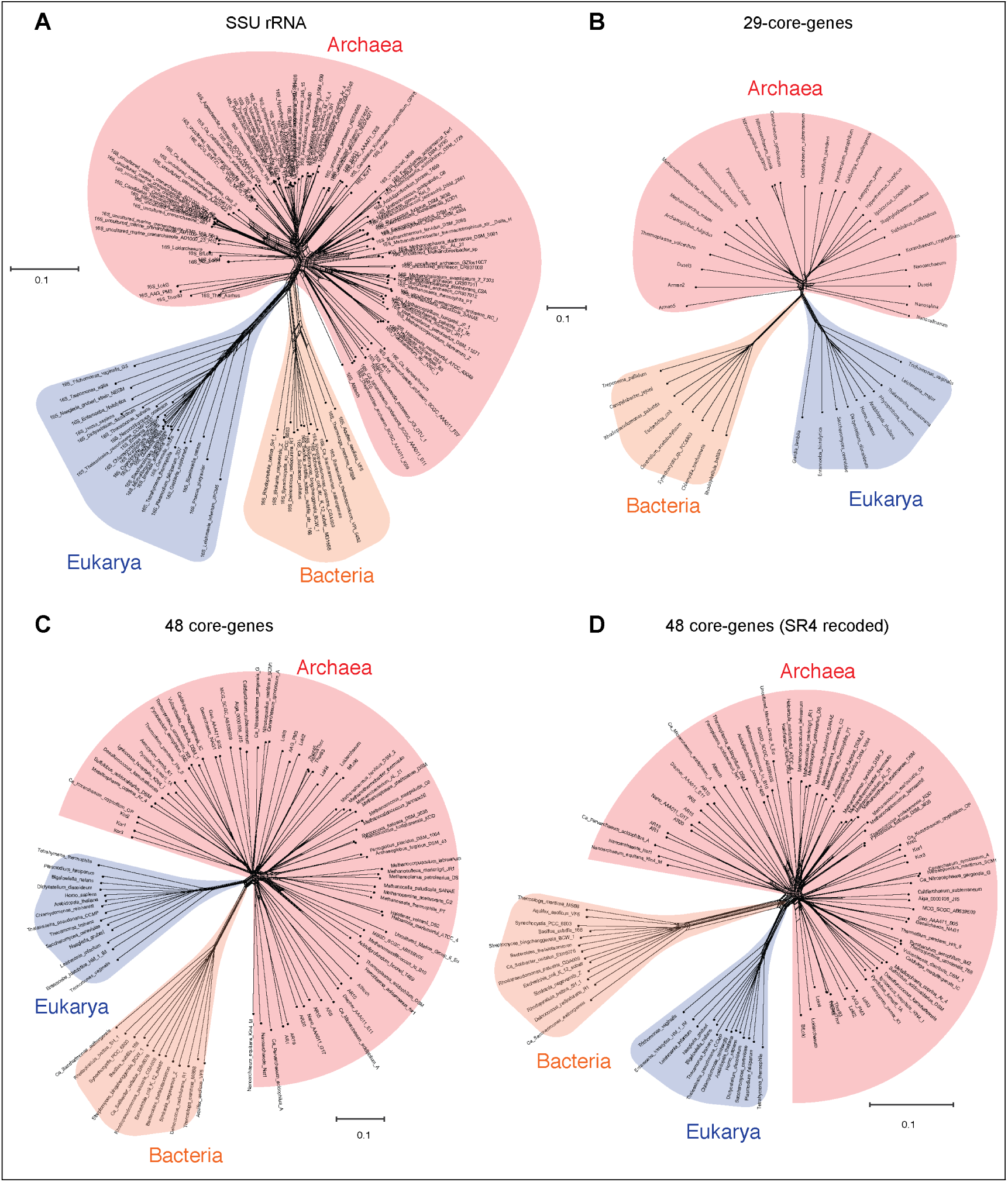
Data-display networks depicting the character conflicts in datasets that employ different character types: nucleotides or amino acids, to resolve the Tree of Life. (A) SSU rRNA alignment of 1,462 characters. Concatenated protein sequence alignment of (B) 29 core-genes, 8,563 characters; (C) 48 core-genes, 9,868 characters and (D) also 48 core-genes, 9,868 SR4 recoded characters (data simplified from 20 to 4 character-states). Each network is constructed from a neighbor-net analysis based on the observed genetic distance (p-distance) and displayed as an equal angle split network. Edge (branch) lengths correspond to the support for character bipartitions (splits), and reticulations in the tree correspond to character conflicts. Datasets in (A), (C) and (D) are from Ref. 18, and in (B) is from Ref. 16.

Likewise, the concatenated protein sequence alignment of the so-called ‘genealogy defining core of genes’ (24) – a set of conserved single-copy genes – also does not support a unique archael lineage. Fig. 1B is a DDN derived from a neighbor-net analysis of 8,563 characters in 29 concatenated core-genes (16), while Fig. 1C,D is based on 9,868 characters in 44 concatenated core-genes (also from (18)). Even taken together, none of the standard marker gene datasets are likely to support the monophyly of the Archaea — a key assertion of the three-domains hypothesis (25). Simply put, there is not enough information in the core-gene datasets to resolve the archaeal radiation, or to determine whether the Archaea are really unique compared to the Bacteria and Eukarya. However, other complex features — including molecular, biochemical and phenotypic characters, as well as ecological adaptations — support the uniqueness of the Archaea. These idiosyncratic archaeal characters include the subunit composition of supramolecular complexes like the ribosome, DNA- and RNA-polymerases, biochemical composition of cell membranes, cell walls, and physiological adaptations to energy-starved environments, among other things (26, 27).

### Complex phylogenomic characters minimize uncertainties regarding the uniqueness of the Archaea

A nucleotide is the smallest possible locus, and an amino acid is a proxy for a locus of a nucleotide triplet. Unlike the elementary amino acid- or nucleotide-characters in the core-genes dataset (Fig.1), the DDN in Fig. 2 is based on complex molecular characters: large genomic loci that are formed by distinct permutations of elementary characters. In this case the loci correspond to protein-domains, typically ∼200 amino acids (600 nucleotides) long. Each protein-domain is unique: with a distinct sequence profile, three-dimensional (3D) structure and function (Fig. 3). Neighbor-net analysis of protein-domain data coded as binary characters (presence/absence) is based on the Hamming distance (identical to the p-distance used in Fig.1). Here the Archaea also form a distinct well-supported cluster, as do the Bacteria and the Eukarya.

**Figure 2.**
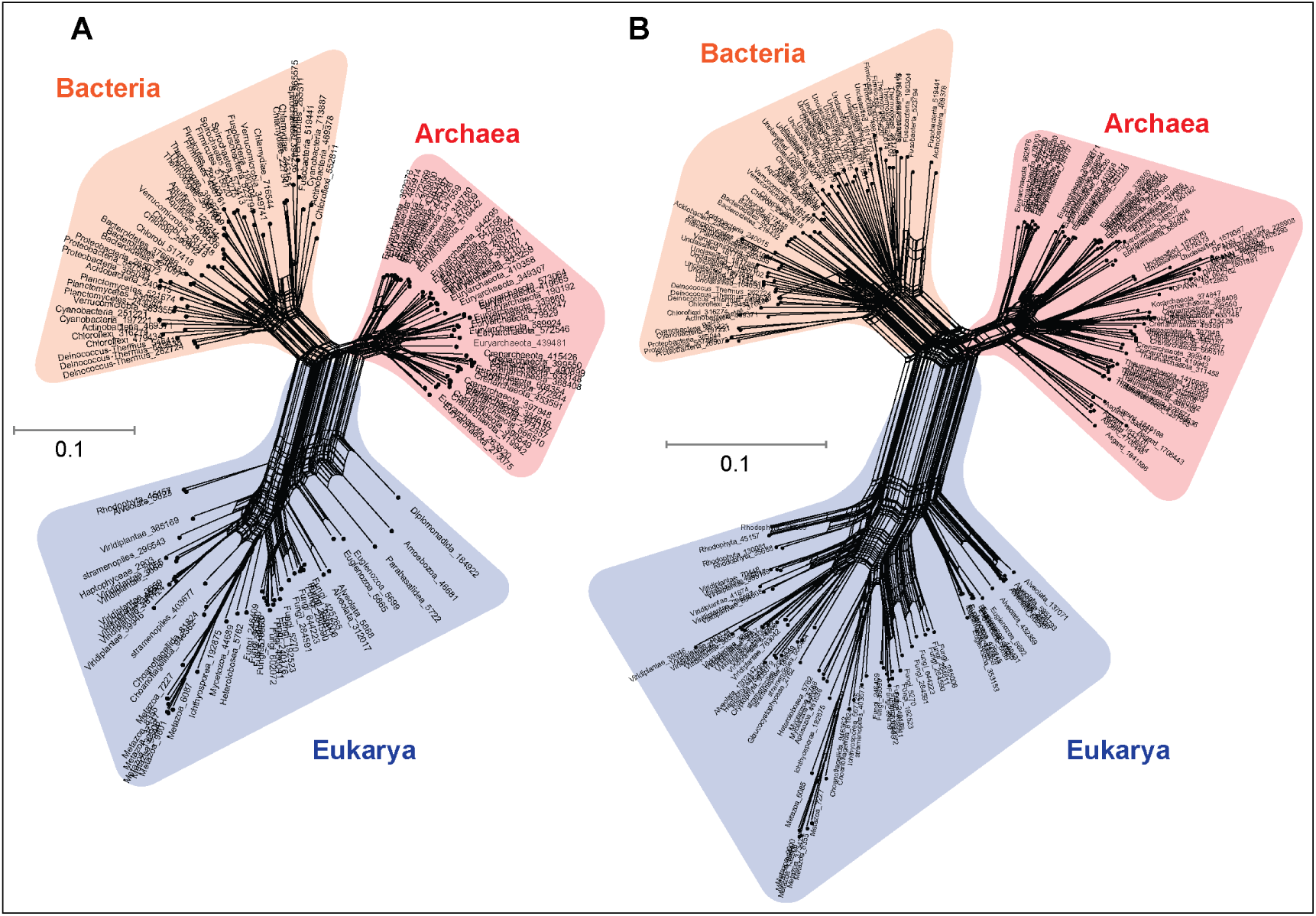
Data-display networks (DDN) depicting character conflicts among complex molecular characters: large genomic loci corresponding to protein-domains as opposed to elementary characters (individual nucleotides or amino acids). (A) Neighbor-net analysis based on Hamming distance (identical to the p-distance used in Fig.1) of 1,732 characters sampled from 141 species. (B) DDN based on an enriched taxon sampling of 81 additional species totaling 222 species and a modest increase to 1,738 characters. The dataset in (A) is from Ref. 9, which was updated with novel species to represent the recently described archaeal and bacterial species (4, 11, 18).

**Figure 3.**
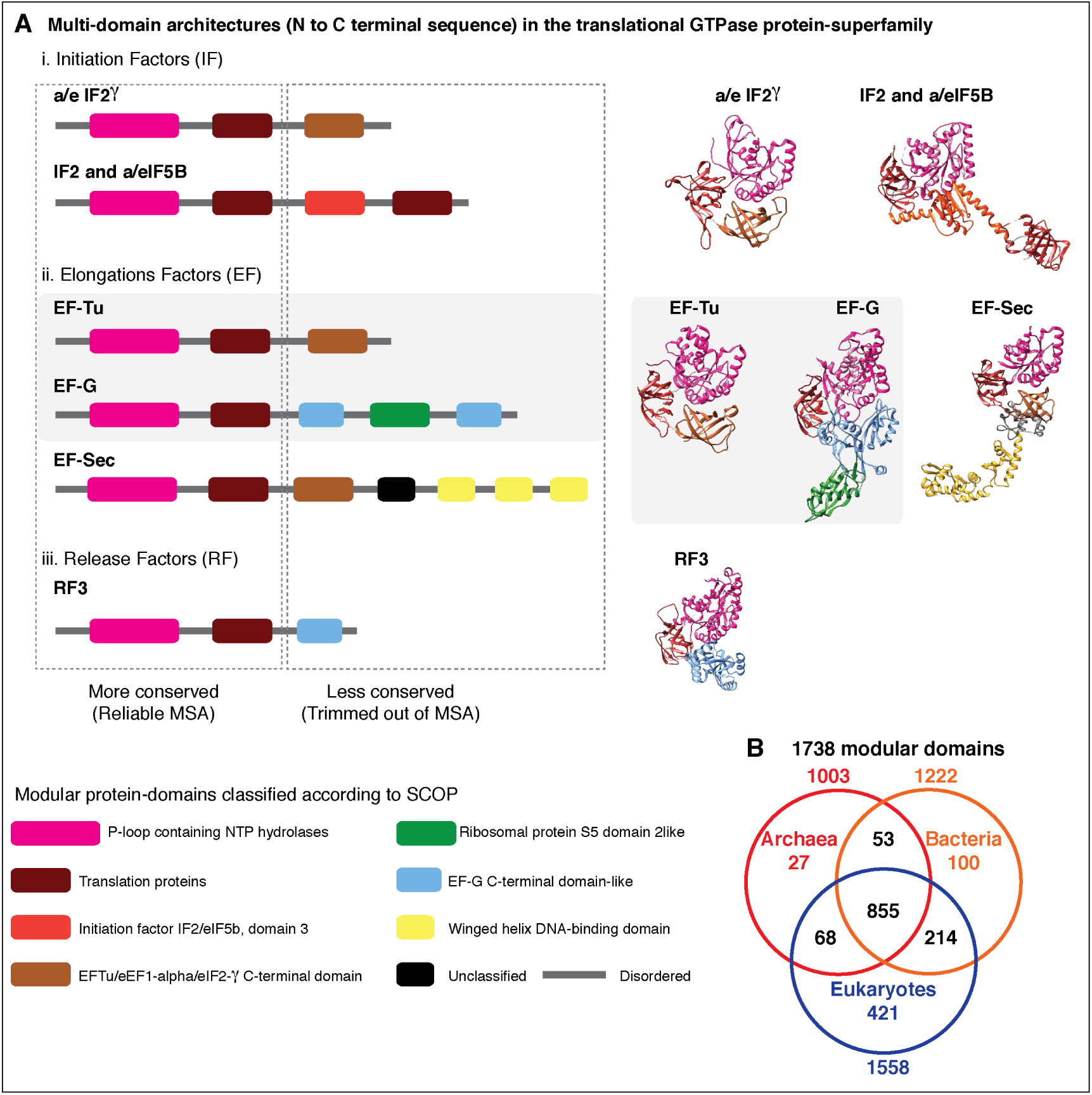
Alignment uncertainty in closely related proteins due to domain recombination. (A) Multi-domain architecture (MDA) of the translational GTPase superfamily based on recombination of 8 modular domains. 57 distinct families with varying MDAs are known, of which 6 canonical families are shown as a schematic on the left and the corresponding 3D folds on the right. Amino acid sequences of only 2 of the 8 conserved domains can be aligned with confidence for use in phylogenetic analysis. The length of the alignment varies from 200-300 amino acids depending on the sequence diversity sampled (13, 32). The EF-Tu—EF-G paralogous pair employed as pseudo-outgroups for the classical rooting of the rRNA tree is highlighted. (B) Phyletic distribution of 1,738 out the 2,000 distinct SCOP-domains sampled from 222 species used for phylogenetic analyses in the present study. About 70 percent of the domains are widely distributed across the sampled taxonomic diversity.

Fig 2A is a DDN based on the dataset that includes protein-domain cohorts of 141 species, used in a phylogenomic analysis to resolve the uncertainties at the root of the ToL (28). Compared to the data in Fig. 1, the taxonomic diversity sampled for the Bacteria and Eukarya is more extensive, but less extensive for the Archaea; it is composed of the traditional groups Euryarchaeota and Crenarchaeota. Fig. 2B is a DDN of an enriched sampling of 81 additional species, which includes representatives of the newly described archaeal groups: TACK (29), DPANN (4), and Asgard group including the Lokiarchaeota (18). In addition, species sampling was enhanced with representatives from the candidate phyla described for Bacteria, and with unicellular species of Eukarya. The complete list of species analyzed is in SI Table 1.

Notably, the extension of the protein-domain cohort was insignificant, from 1,732 to 1,738 distinct domains (characters). Based on the well-supported splits in the DDN that form a distinct archaeal cluster, the Archaea are likely to be a monophyletic group (clade) in phylogenies inferred from these datasets.

### Employing complex molecular characters maximizes representation of orthologous, non-recombining genomic loci, and thus phylogenetic signal

Despite the superficial similarity of the DDNs in Fig.1 and Fig.2, they are both qualitatively and quantitatively different codings of genome sequences. As opposed to tracing the history of at most 50 loci in the standard core-genes datasets (Fig. 1), up to 60 fold (1738 loci) more information can be represented when genome sequences are coded as protein-domain characters (Fig. 2). Currently 2,000 unique domains are described by SCOP (Structural Classification of Proteins) (30). The phyletic distribution of 1,738 domains identified in the 222 representative species sampled here is shown in a Venn diagram (Fig. 3B).

Genomic loci that can be aligned with high confidence using MSA algorithms are typically more conserved than those loci for which alignment uncertainty is high. Such ambiguously aligned regions of sequences are routinely trimmed off before phylogenetic analyses (31). Typically, the conserved well-aligned regions correspond to protein domains with highly ordered three-dimensional (3D) structures with specific 3D folds (Fig. 3A). A closer look at the 29 core-genes dataset shows that the concatenated-MSA corresponds to a total of 27 distinct protein domains or genomic loci (Table 1).

**Table 1.**
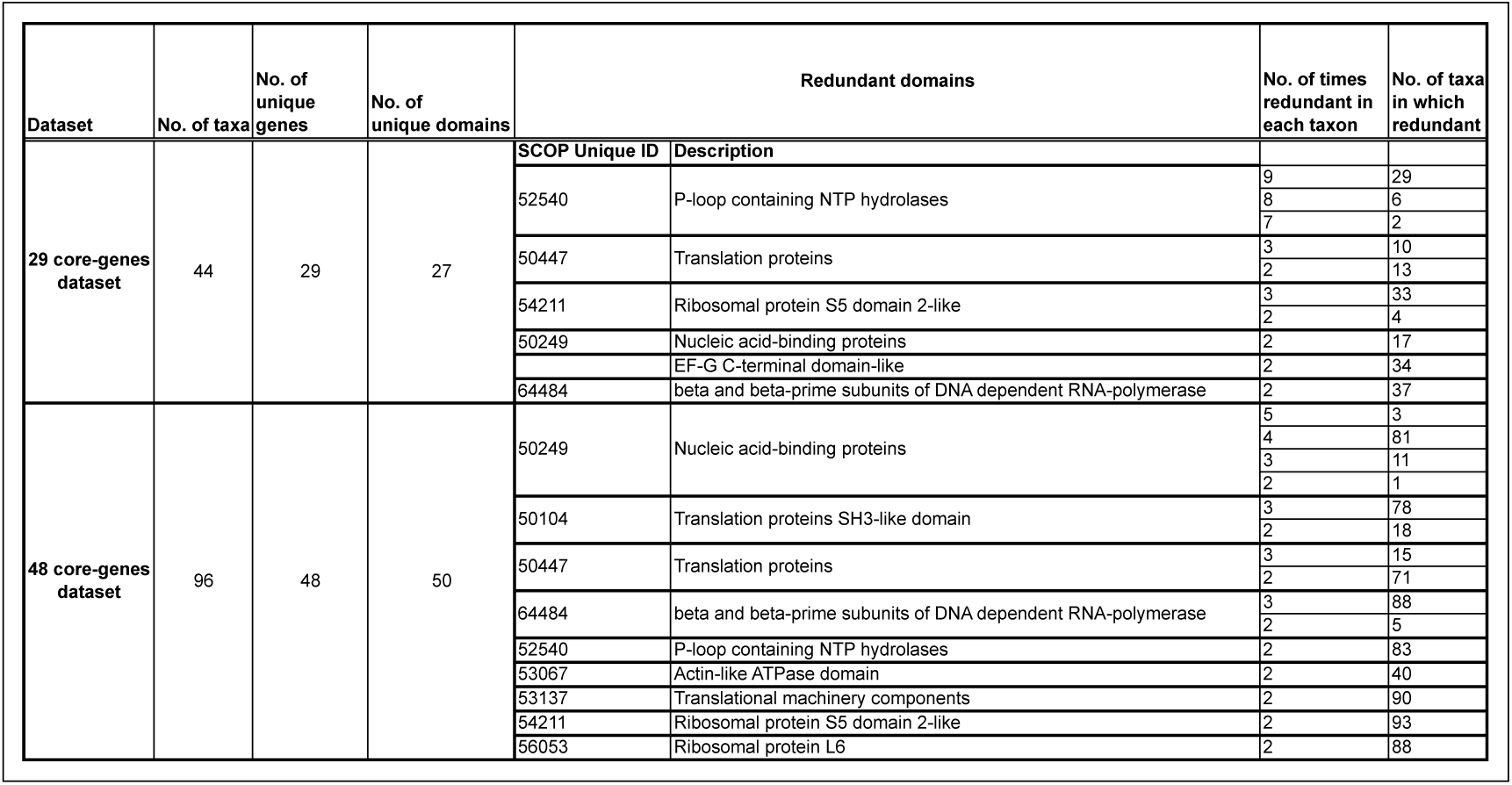
Redundant representation of protein-domains in concatenated core-genes datasets. The P-loop NTP hydrolase domain is one of the most prevalent domains. Genomic loci coding for P-loop hydrolase domain are represented 8-9 times in each species in the single-copy genes employed from core-genes multiple sequence alignments. Redundant loci in the core-genes datasets vary depending on the genes and species sampled for phylogenomic analyses.

The number of loci sampled from different species varies between 20 and 27, since not all loci are found in all species. While some loci are absent in some species, some loci are redundant. For instance, the P-loop NTP hydrolase domain, one of the most prevalent protein domains, is represented up to 9 times in many species (Table 1). Many central cellular functions are driven by the conformational changes in proteins induced by the hydrolysis of nucleoside triphosphate (NTP) catalyzed by the P-loop domain. Out of a total of 27 distinct domains, 7 are redundant, with two or more copies represented per species. Similarly, 9 of the 50 domains have a redundant representation in the 44 core-gene dataset (Table 1). The observed redundancy of the genomic loci in the core-genes alignments is inconsistent with the common (and typically untested) assumption of using single-copy genes as a proxy for orthologous loci sampled for phylogenetic analysis. In contrast, the protein-domain datasets are composed of unique loci (Fig. 3B).

Regions of sequences that are trimmed usually show higher variability in length, are less ordered and are known to accumulate insertion and deletion (indel) mutations at a higher frequency than in the regions that correspond to folded domains (33). These variable, structurally disordered regions, which flank the structurally ordered domains, link different domains in multi-domain proteins (Fig. 3A). Multi-domain architecture (MDA), the N-to-C terminal sequence of domain arrangement, is distinct for a protein family, and differs in closely related protein families with similar functions (Fig. 3A). The variation in MDA also relates to alignment uncertainties.

### Data quality affects model complexity required to explain phylogenetic datasets

Resolving the paraphyly or monophyly of the Archaea is relevant to determining whether the Eocyte tree (Fig. 4A) or the Three-domains tree (Fig. 4B), respectively, is a better-supported hypothesis. Recovering the Eocyte tree typically requires implementing complex models of sequence evolution rather than their relatively simpler versions (10). In general, complex models tend to fit the data better. For instance, according to a model selection test for the 29 core-genes dataset, the LG model of protein sequence evolution is a better-fitting model than other standard models, such as the WAG or JTT substitution model (SI-Table 2), as reported previously (16). Further, a relatively more complex version of the LG model, with multiple rate-categories is a better-fitting model than the simpler single-rate-category model, as seen from the higher likelihoods (Fig. 4C; SI-Table 2).

**Figure 4.**
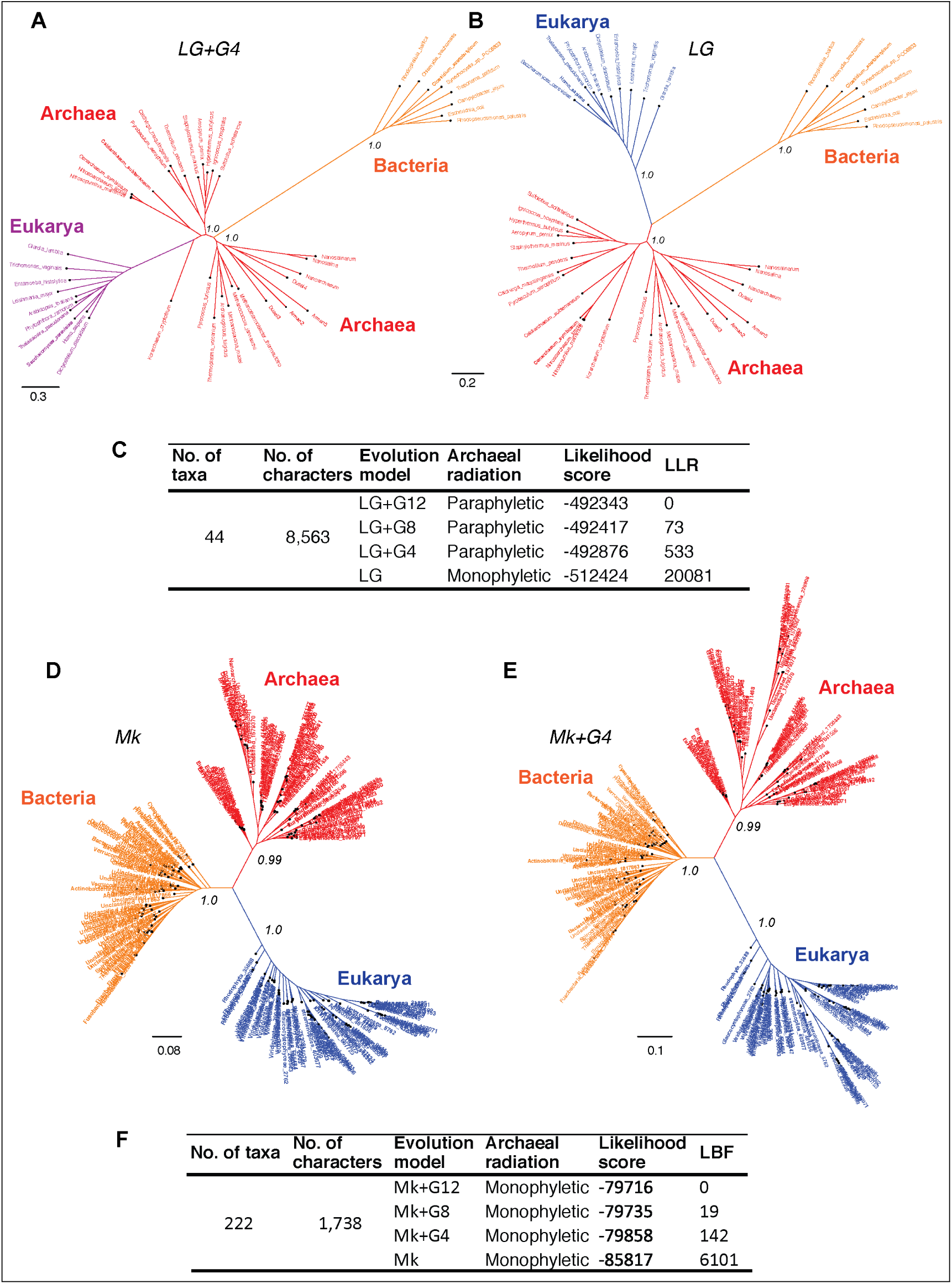
Comparison of concatenated-gene trees derived from amino acid characters and genome trees derived from protein-domain characters. Branch support is shown only for the major branches. Scale bars represent the expected number of changes per character. (A), (B) Core-genes-tree derived from a better-fitting model (LG+G4) and a worse fitting mode (LG), respectively, of amino acid substitutions. (C) Model fit to data is ranked according the log likelihood ratio (LLR) scores. LLR scores are computed as the difference from the best-fitting model (LG+G12) of the likelihood scores estimated in PhyML. Thus, larger LLR values indicate less support for that model/tree relative to the most-likely model/tree. Substitution rate heterogeneity is approximated with 4, 8 or 12 rate categories in the complex models, but with a single rate category in the simpler model. (D), (E) are genome-trees derived from a better-fitting model (Mk+G4) and a worse fitting model (Mk), respectively, of protein-domain innovation. (F) Model fit to data is ranked according log Bayes factor (LBF) scores, which like LLR scores are the log odds of the hypotheses. LBF scores are computed as the difference in likelihood scores estimated in MrBayes.

A complex, multiple rate-categories model accounts for site-specific substitution rate variation. Substitution-rate heterogeneity across different sites in the multiple-sequence alignment (MSA) was approximated using a discrete Gamma model with 4, 8 or 12 rate categories (LG+G4, LG+G8 or LG+G12, respectively). The Archaea is consistent with a paraphyletic group in trees derived from the rate-heterogeneous versions of the LG model (Fig. 4A). Furthermore, the fit of the data improves with the increase in complexity of the substitution model (Fig. 4C). Model complexity increases with any increase in the number of rate categories and/or the associated numbers of parameters that need to be estimated. However, with a relatively simpler version, in which the substitution-rates are approximated to a single rate-category (i.e. a rate-homogeneous LG model), the Archaea are consistent with a monophyletic group (Fig. 4B).

In contrast, trees inferred from the protein-domain datasets are consistent with monophyly of the Archaea irrespective of the complexity of the underlying model (Fig. 4D-F). The Mk model (Markov k model) is the best-known probabilistic model of discrete character evolution, particularly of complex characters coded as binary-state characters (35, 36). Since the Mk model assumes a stochastic process of evolution, it is able to estimate multiple state changes along the same branch. Implementing a simpler rate-homogeneous version of the Mk model (Fig. 4D), as well as more complex rate-heterogeneous versions with 4, 8 or 12 rate categories (Mk+G4, Mk+G8 or Mk+G12, respectively), also recovered trees that are consistent with the monophyly of the Archaea (Fig. 4E). The tree derived from the Mk+G4 model is shown in Fig. 4E. While the tree derived from Mk+G8 model is identical to the Mk+G4 tree, the Mk+G12 tree is almost identical with minor differences in the bacterial sub-groups (SI-Fig. 1)

In all cases, bipartitions for Archaea show strong support with posterior probability (PP) of 0.99 while that of Bacteria and Eukarya is supported with a PP of 1.0: in spite of substantially different fits of the data. The uniqueness of the Archaea is almost unambiguous in this case (but see next section).

### Siblings and cousins are indistinguishable when reversible models are employed

Although a DDN is useful to identify and diagnose character conflicts in phylogenetic datasets and to postulate evolutionary hypotheses, a DDN by itself cannot be interpreted as an evolutionary network, because the edges do not necessarily represent evolutionary phenomena and the nodes do not represent ancestors (22, 23). Therefore, evolutionary relationships cannot be inferred from a DDN. Likewise, evolutionary relationships cannot be inferred from unrooted trees, even though nodes in an unrooted tree do represent ancestors and an evolution model defines the branches (see Fig. 5A).

**Figure 5.**
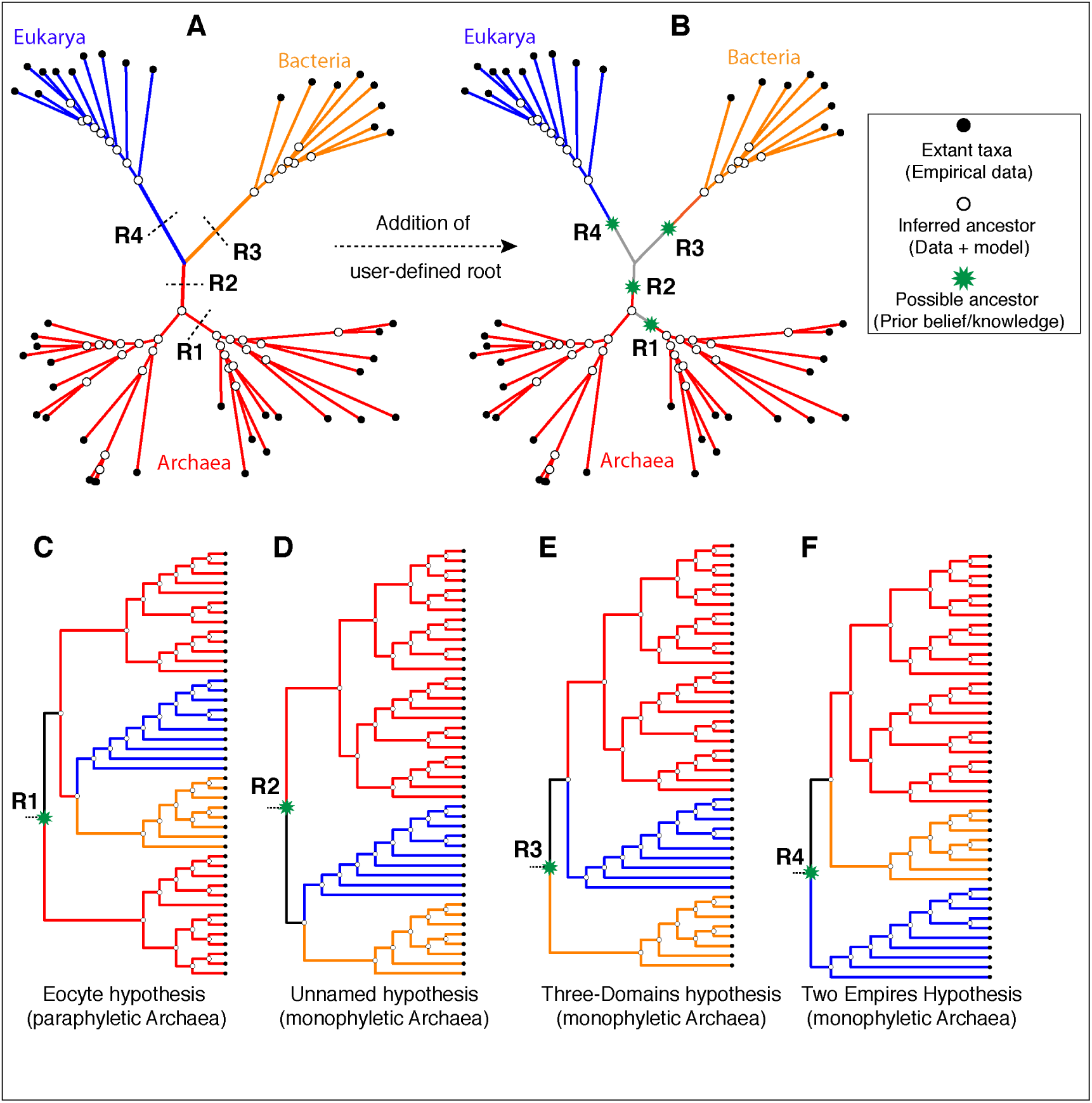
Effect of alternative *ad hoc* rootings on the phylogenetic classification of archaeal biodiversity. (A) An unrooted tree is not fully resolved into bipartitions at the root of the tree (i.e. a polytomous rather than a dichotomous root branching) and thus precludes identification of sister group relationships. It is common practice to add a user-specified root *a posteriori* based on prior knowledge (or belief) of the investigator. Four possible (of many) rootings R1-R4 are shown. (B) Operationally, adding a root (rooting) *a posteriori* amounts to adding new information – a new bipartition and an ancestor as well as an evolutionary polarity – that is independent of the source data. (C-F) The different possible evolutionary relationships of the Archaea to other taxa, depending on the position of the root, are shown. Rooting is necessary to determine the recency of common ancestry as well the temporal order of key evolutionary transitions that define phylogenetic relationships.

Given that a primary objective of phylogenetic analyses is to identify clades and the relationships between these clades, it is not possible to interpret an unrooted tree meaningfully without rooting the tree (see Fig. 5A). Identifying the root is essential to: (i) distinguish between ancestral and derived states of characters, (ii) determine the ancestor-descendant polarity of taxa, and (iii) diagnose clades and sister-group relationships (Fig. 5). Yet, most phylogenetic software construct only unrooted trees, which are then consistent with several rooted trees (Fig. 5 C-F). Thus, an unrooted tree, unlike a rooted tree, is not an evolutionary (phylogenetic) tree *per se*, since it is a minimally defined hypothesis of evolution or of relationships; it is, nevertheless, useful to rule out many possible bipartitions and groups (34, 35). However, an unrooted tree cannot be fully resolved into bipartitions, because an unresolved polytomy (a trichotomy in this case) exists near the root of the tree (Fig. 5A), which otherwise corresponds to the deepest split (root) in a rooted tree (Fig. 5, C-F).

Resolving the polytomy requires identifying the root of the tree. The identity of the root corresponds, in principle, to any one of the possible ancestors as follows:

i. Any one of the inferred-ancestors at the resolved bipartitions (open circles in Fig. 5A), or
ii. Any one of the yet-to-be-inferred-ancestors that lies along the stem-branches of the unresolved polytomy (dashed lines in Fig. 5A) or along the internal-braches.

In the latter case, rooting the tree *a posteriori* on any of the branches amounts to inserting an additional bipartition and an ancestor that is neither inferred from the source data nor deduced from the underlying character evolution model. Since standard evolution models employed routinely cannot resolve the polytomy, rooting, and hence interpreting the Tree of Life depends on:

i. Prior knowledge — eg., fossils or a known sister-group (outgroup), or
ii. Prior beliefs/expectations of the investigators — eg., simple is primitive (37, 38), bacteria are primitive (39, 40), archaea are primitive (1), etc.

Both of these options are independent of the data used to infer the unrooted ToL. Some possible rootings and the resulting rooted-tree topologies are shown as cladograms in Fig. 5, C-F. If the root lies on any of the internal branches (e.g. R1 in Fig. 5,A-C), or corresponds to one of the internal nodes, within the archael radiation, the Archaea would not constitute a unique clade (Fig. 5C). However, if the root lies on one of the stem-branches (R2/R3/R4 in Fig. 5 A, B), monophyly of the Archaea would be unambiguous (Fig. 5 D-F). Determining the evolutionary relationship of the Archaea to other taxa, though, requires identifying the root.

Directional evolution models, unlike reversible models, are able to identify the polarity of state transitions, and thus the root of a tree (41-43). Therefore, the uncertainty due to a polytomous root branching is not an issue (Fig 6A). Moreover, directional evolution models are useful to evaluate the empirical support for prior beliefs about the universal common ancestor (UCA) at the root of the ToL (28). A Bayesian model selection test implemented to detect directional trends (43) chooses the directional model, overwhelmingly (Fig. 6B), over the unpolarized model for the protein-domain dataset in Fig. 2B, as reported previously for the dataset in Fig. 2A (28). Further, the best-supported rooting corresponds to root R4 (Fig. 5F and Fig. 6A) — monophyly of the Archaea is maximally supported (PP of 1.0). Furthermore, the sister-group relationship of the Archaea to the Bacteria is maximally supported (PP 1.0). Accordingly, a higher order taxon, Akaryotes, proposed earlier (44) forms a well-supported clade. Thus Akaryotes (or Akarya) and Eukarya are sister clades that diverge from the UCA at the root of the ToL, also as reported previously (28).

**Figure 6.**
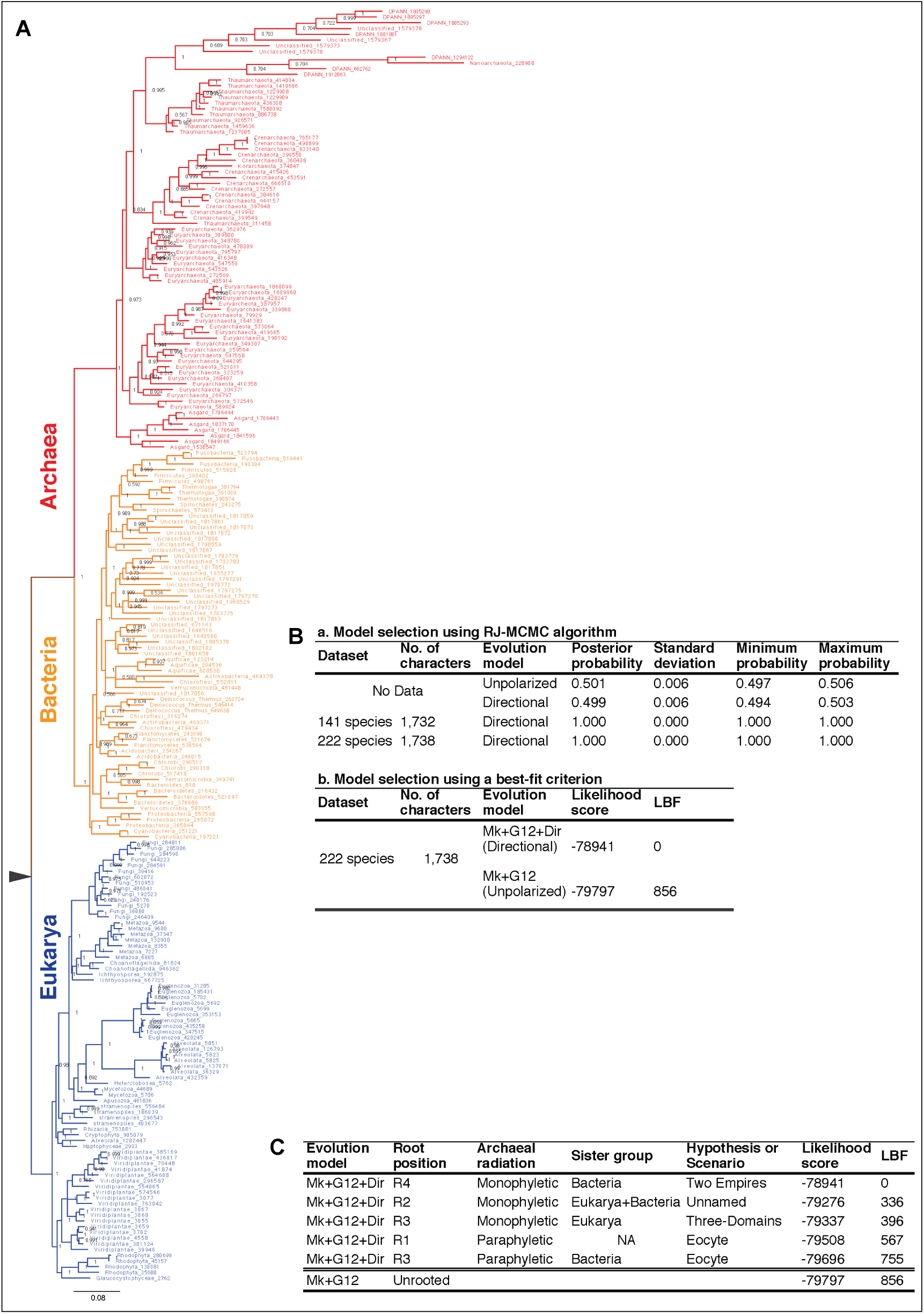
(A) Rooted tree of life inferred from patterns of inheritance of functional genomic-signatures. A dichotomous classification of the diversity of life such that Archaea is a sister group to Bacteria, which together constitute a clade of akaryotes (Akarya). Eukarya and Akarya are sister-clades that diverge from the root of the tree of life. Each clade is supported by the highest posterior probability of 1.0. The phylogeny supports a scenario of independent origins and descent of eukaryotes and akaryotes. (B) Model selection tests identify, overwhelmingly, directional evolution models to be better-fitting models. (C) Alternative rootings, and accordingly alternative classifications or scenarios for the origins of the major clades of life, are much less probable and not supported.

Alternative rootings are much less likely, and are not supported (Fig. 6C). Accordingly, independent origin of the eukaryotes as well akaryotes is the best-supported scenario. The Three-domains tree (root R3, Fig. 5E) is 10171 times less likely, and the scenario proposed by the Eocyte hypothesis (root R1, Fig. 5C) is highly unlikely. The common belief that simple is primitive, as well as beliefs that archaea are primitive or that archaea and bacteria evolved before eukaryotes, are not supported either.

## Discussion

### Improving data quality can be more effective for resolving recalcitrant branches than increasing model complexity

In the phylogenetic literature, the concept of data quality refers to the quality or the strength of the phylogenetic signal that can be extracted from the data. The strength of the phylogenetic signal is proportional to the confidence with which unique state-transitions can be determined for a given set of characters on a given tree. Ideally, historically unique character transitions that entail rare evolutionary innovations are desirable, to identify patterns of uniquely shared innovations (synapomorphies) among lineages. Synapomorphies are the diagnostic features used for assessing lineage-specific inheritance of evolutionary innovations. Therefore identifying character transitions that are likely to be low probability events is a basic requirement for the accuracy of phylogenetic analysis.

In their pioneering studies, Woese and colleagues identified unique features of the SSU rRNA s– [oligonucleotide] “signatures” – that were six nucleotides or longer, to determine evolutionary relationships (2). An underlying assumption was that the probability of occurrence of the same set of oligomer signatures by chance, in non-homologous sequences, is low in a large molecule like SSU rRNA (1500-2000 nucleotides). Oligomers shorter than six nucleotides were statistically less likely to be efficient markers of homology (45). Thus SSU rRNA was an information-rich molecule to identify homologous signatures (characters) useful for phylogenetic analysis.

However, as sequencing of full-length rRNAs and statistical models of nucleotide substitution became common, complex oligomer-characters were replaced by elementary nucleotide-characters; and more recently by amino acid characters. Identifying rare or historically unique substitutions in empirical datasets has proven to be difficult (46, 47), consequently the uncertainty of resolving the deeper branches of the Tree of Life using marker-gene sequences remains high. A primary reason is the relatively higher prevalence of phylogenetic noise (homoplasy) in primary sequence datasets (Fig. 1), due to the characteristic redundancy of nucleotide and amino acid substitutions and the resulting difficulty in distinguishing phylogenetic noise from signal (homology) (48, 49). Better-fitting (or best-fitting) models are expected to extract phylogenetic signal more efficiently and thus explain the data better, but tend to be more complex than worse-fitting models (Fig. 4 C, F). Increasingly sophisticated statistical models that have been developed over the years have only marginally improved the situation (50, 51). Although increasing model complexity can correct errors of estimation and improve the fit of the data to the tree, it is not a solution to improve phylogenetic signal, especially when not present in the source data.

The idea of ‘oligonucleotide-signatures’ used for estimating a gene phylogeny has been extended, naturally, to infer a genome phylogeny (52). The signatures were defined in terms of protein-coding genes that were shared among the Archaea. However, as proteins are mosaics of domains, domains are unique genomic signatures (Fig. 3). Protein domains defined by SCOP correspond to complex ‘multi-dimensional signatures’ defined by: (i) a unique 3D fold, (ii) a distinct sequence profile, and a characteristic function. Though domain recombination is frequent, substitution of one protein domain for another has not been observed in homologous proteins (Fig. 3). For phylogenomic applications, protein domains are ‘sequence signatures’ that essentially correspond to single-copy orthologous loci when coded as binary-state characters (presence/absence). These sequence signatures are consistent with unique, non-recombining genomic loci, and are identified using sophisticated statistical models — profile hidden Markov models (pHMMs) (53, 54) — that can be used routinely to annotate and curate genome sequences in automated pipelines (55, 56).

Character recoding is found to be effective in reducing the noise/redundancy in the data, and thus uncertainties in phylogenetic reconstructions (57). This is a form of data simplification wherein the number of amino acid alphabets is reduced to a smaller set of alphabets that are frequently substituted for each other, usually reduced from 20 to 6. Character recoding into reduced alphabets is useful in cases were compositional heterogeneity or substitution saturation is high (57). However, datasets in which phylogenetic noise is inherently limited are more desirable, to minimize ambiguities. Like amino acids, protein domains are also modular alphabets, albeit higher order and more complex alphabets of proteins. Moreover, unlike the 20 standard amino acids, there are approximately 2,000 unique protein domains identified at present according to SCOP (33). The number is expected to increase; the theoretical estimates range between 4,000 and 10,000 distinct domain modules, depending on the classification scheme (58). Coding features as binary characters is the simplest possible representation of data for describing historically unique events. For these reasons, protein domains are highly effective phylogenetic markers for which character-homology can be validated through more than one property, statistically significant (i) sequence similarity, (ii) 3D structure similarity; and (iii) function similarity. In addition, employing genomic loci for protein domains maximizes the genomic information that can be employed for phylogenetic analysis. Even though many other genomic features are known to be useful markers (59), protein domains are the most conserved as well as most widely applicable genomic characters (Fig. 3B).

### Sorting vertical evolution (signal) and horizontal evolution (noise)

Single-copy genes are employed as phylogenetic markers to minimize phylogenetic noise caused by reticulate evolution, including hybridization, introgression, recombination, horizontal transfer (HT), duplication-loss (DL), or incomplete lineage sorting (ILS) of genomic loci. However, the noise observed in the DDNs based on MSA of core-genes (Fig. 1) cannot be directly related to any of the above genome-scale reticulations, since the characters are individual nucleotides or amino acids. Apart from stochastic character conflicts, the observed conflicts are better explained by convergent substitutions, given the redundancy of substitutions. Convergent substitutions caused either due to stringent selection or by chance are a well-recognized form of homoplasy in gene-sequence data (46, 49, 60), and based on recent genome-scale analyses it is now known to be rampant (61, 62).

The observed noise in the DDNs based on protein-domain characters (Fig. 2), however, can be related directly to genome-scale reticulation processes and homoplasies. In general, homoplasy implies evolutionary convergence, parallelism or character reversals caused by multiple processes. In contrast, homology implies only one process: inheritance of traits that evolved in the common ancestor and were passed to its descendants. Operationally, tree-based assessment of homology requires tracing the phylogenetic continuity of characters (and states), whereas homoplasy manifests as discontinuities along the tree. Since clades are diagnosed on the basis of shared innovations (synapomorphies) and defined by ancestry (63, 64), accuracy of a phylogeny depends on an accurate assessment of homology — unambiguous identification of relative synapomorphies on a best fitting tree.

Identifying homoplasies caused by character reversals, i.e. reversal to ancestral states requires identification of the ancestral state of the characters under study. However, implementing reversible models precludes the estimation of ancestral states, in the absence of sister groups (outgroups) or other external references. Thus, the critical distinction between shared ancestral homology (symplesiomorphy) and shared derived homology (synapomorphy) is not possible with unrooted trees derived from standard reversible models. Hence, unrooted trees (Fig. 4) are not evolutionary (phylogenetic) trees *per se*, as they are uninformative about the evolutionary polarity (65, 66). Thus, identifying the root (or root-state) is crucial to (i) determine the polarity of state transitions, (ii) identify synapomorphies, and (iii) diagnose clades.

For complex characters such as protein domains, character homology can be determined with high confidence using sophisticated statistical models (HMMs). Homology of a protein domain implies that the *de novo* evolution of a genomic locus corresponding to that protein domain is a unique historical event. Therefore, homoplasy due to convergences and parallelisms is highly improbable (67, 68). Although a handful of cases of convergent evolution of 3D structures is known, these instances relate to relatively simple 3D folds coded for by relatively simple sequence repeats (69). However, the vast majority of domains identified by SCOP correspond to polypeptides that are on average 200 residues long with unique sequence profiles (55, 67). Thus, identifying homoplasy in the protein-domain datasets depends largely on estimating reversals, which will be cases of secondary gains/losses; for instance gain-loss-regain events caused by HT or DL-HT. Such secondary gains are more likely to correspond to HT events than to convergent evolution, for reasons specified above. Instances of reversals are minimal, as seen from the strong directional trends detected in the data (Fig. 6B). Thus, uncertainty in determining polarity of state transitions, identifying synapomorphies and diagnosing clades is minimized.

Moreover, because clades are associated with the emergence and inheritance of evolutionary novelties, the discovery of clades is fundamental for describing and diagnosing sister group differences, which is a primary objective of modern systematics (70). A well-recognized deficiency of phylogenetic inference based on primary sequences is the abstraction of evolutionary ‘information’ (52), often into less tangible quantitative measures. For instance, ‘information’ relevant to diagnosing clades and support for clades is abstracted to branch lengths. Branch-length estimation is, ideally, a function of the source data and the underlying model. However, in the core-genes dataset the estimated branch lengths and the resulting tree is an expression of the model rather than of the data (Fig. 4 A, B). Some pertinent questions then are: should diagnosis of clades and the features by which clades are identified be delegated to, and restricted to, substitution mutations in a small set of loci and substitution models? Are substitution mutations in 40-50 loci more informative, or the birth and death of unique genomic loci more informative?

Proponents of the total evidence approach recommend that all relevant information — molecular, biochemical, anatomical, morphological, fossils — should be used to reconstruct evolutionary history, yet genome sequences are the most widely applicable data at present (59, 71). Accordingly, phylogenetic classification is, in practice, a classification of genomes. There is no *a priori* theoretical reason that phylogenetic inference should be restricted to a small set of genomic loci corresponding to the core genes, nor is there a reason for limiting phylogenetic models to interpreting patterns of substitution mutations alone. The ease of sequencing and the practical convenience of assembling large character matrices, by themselves, are no longer compelling reasons to adhere to the traditional marker gene analysis. Annotations for reference genomes of homologous protein domains identified by SCOP and other protein-classification schemes, as well as tools for identifying corresponding sequence signatures, are readily available in public databases. An added advantage is that the biochemical function and molecular phenotype of the domains are readily accessible as well, through additional resources including protein data bank (PDB) and InterPro.

### Vertical and horizontal classification

For decades, biologists have been faced with a choice between so-called horizontal (Linnean) and vertical (Darwinian) classification of biodiversity (72). The similarity of both schools of systematics concerns the identification of “signatures” or sets of characteristic features that codify evolutionary relationships (52, 63, 72). But the former emphasizes the unity of contemporary groups, i.e. those at a similar evolutionary state, and therefore separates ancestors from descendants, while the latter emphasizes the unity of the ancestors and separates descendants that diverge from a common ancestry (72). Vertical classification is more consistent with the concept of lineal descent, and is the predominant paradigm for which the operational methodology and the algorithmic logic were laid out as the principles of phylogenetic systematics (63, 73). Accordingly, determining the ancestor-descendant polarity, starting from the universal common ancestor (UCA) at the root of the Tree of Life, is crucial for accurately reconstructing the path of evolutionary descent.

The classical rooting of the (rRNA) ToL based on the EF-Tu—EF-G paralogous pair (74, 75) is known to be error-prone and highly ambiguous, due to LBA artifacts (14, 75). Remarkably, sequences corresponding to only one of the two conserved domains common to EF-Tu and EF-G (200 residues in the P-loop-containing NTP hydrolase domain (Fig. 3A)) can be aligned with confidence (13). Implementing better-fitting substitution models results in two alternative rootings (R1 and R4 in Fig. 5), which relate to distinct, irreconcilable scenarios (13), similar to scenarios in Fig 5C and 5F. Moreover, the EF-Tu—EF-G paralogous pair is only 2 of 57 known paralogs of the translational GTPase protein superfamily (32). Thus the assumption that EF-Tu—EF-G duplication is a unique event, which is essential for the paralogous outgroup-rooting method, is untenable.

In the absence of prior knowledge of outgroups or of fossils, rooting the Tree of Life is arguably one of the most difficult phylogenetic problems. The conventional practice of *a posteriori* rooting, wherein an unrooted tree is converted into a rooted tree by adding an *ad hoc* root, encourages a subjective interpretation of the ToL. For example, the so-called bacterial rooting of the ToL (root R3; Fig. 5) is the preferred rooting hypothesis to interpret the ToL even though that rooting is not well supported (13). Incorrect rooting may lead to profoundly misleading conclusions about evolutionary scenarios and taxonomic affinities, and it appears to be common in phylogenetic studies (76). Perhaps worse yet seems to be the preponderance of subjective *a posteriori* rooting based on untested preconceptions (e.g. (38, 77)) and scenario-driven erection of taxonomic ranks (78) (e.g. (1, 7, 29)).

### Untangling data bias, model bias and investigator bias (prior beliefs)

Phylogenies, and hence the taxonomies and evolutionary scenarios they support, are falsifiable hypotheses. Statistical hypothesis testing is now an integral part of phylogenetic inference, to quantify the empirical evidence in support of the various plausible evolutionary scenarios. However, common statistical models implemented for phylogenomic analyses are limited to modeling variation in patterns of point mutations, particularly substitution mutations. These statistical models are intimately linked to basic concepts of molecular evolution, such as the universal molecular clock (3), the universal chronometer (77), paralogous outgroup rooting (79), etc., which are gene-centric concepts that were developed to study the gene, during the age of the gene. Moreover, these idealized notions originated from the analyses of relatively small single-gene datasets.

Conventional phylogenomics of multi-locus datasets is a direct extension of the concepts and methods developed for single-locus datasets, which rely exclusively on substitution mutations (49). In contrast, the fundamental concepts of phylogenetic theory: homology, synapomorphy, homoplasy, character polarity, etc., even if idealized, are more generally applicable. And, apparently they are better suited for unique and complex genomic characters rather than for redundant, elementary sequence characters, with regards to determining both qualitative as well as statistical consistency of the data and the underlying assumptions.

Phylogenetic theory that was developed to trace the evolutionary history of organismal species, as well as related methods of discrete character analysis for classifying organismal families (63, 80), was adopted, although not entirely, to determine the evolution and classification of gene families (1, 3). The discovery and initial description of the Archaea was based on the comparative analysis of a single-gene (rRNA) family. However, in spite of the large number of characters that can be analyzed, neither the rRNA genes nor multi-gene concatenations of core-genes have proved to be efficient phylogenetic markers to reliably resolve the evolutionary history and phylogenetic affinities of the Archaea (81, 82).

Uncertainties and errors in phylogenetic inference are primarily errors in adequately distinguishing homologous similarities from homoplastic similarities (49, 65, 83). Homologies, synapomorphies and homoplasies are qualitative inferences, yet are inherently statistical (probabilistic). The probabilistic framework (maximum likelihood and Bayesian methods) has proven to be powerful for quantifying uncertainties and testing alternative hypotheses. Log odds ratios, such as LLR and LBF, are measures of how one changes belief in a hypothesis in light of new evidence (84). Accordingly, directional evolution models are more optimal explanations of the observed distribution of genomic-characters, and such directional trends overwhelmingly support the monophyly of the Archaea, as well as the sisterhood of the Archaea and the Bacteria, i.e. monophyly of Akarya (Fig 6). Data quality is at least as important as the evolution models that are posited to explain the data. Although sophisticated statistical tests for evaluating tree robustness, and for selecting character-evolution models, are becoming a standard feature of phylogenetic software tests for character evaluation are not common. Routines for collecting and curating data upstream of phylogenetic analyses are rather eclectic. Regardless, exclusive reliance on a single data type to resolve different phylogenetic problems might not be practical. Besides, it is an open question as to whether qualitatively different datasets (as in Fig.1 and Fig.2) can be compared effectively. Therefore, polyphasic analyses of different data types that are informative at different phylogenetic depths could be useful, rather than combined analysis. Nevertheless, employing DDNs and other tools of exploratory data analysis could be useful to identify conflicts that arise due to data collection and/or curation errors (22, 23).

## Conclusions

The Tree of Life is primarily a phylogenetic classification that is invaluable to organize and to describe the evolution of biodiversity, explicated through evolutionary scenarios. Phylogenies are hypotheses that mostly relate to extinct ancestors, while taxonomies are hypotheses that largely relate to extant species. Extant species contain distinct combinatorial mosaics of ancestral features (plesiomorphies) and evolutionary novelties (apomorphies). It is remarkable that the uniqueness of the Archaea was identified by the comparative analyses of oligonucleotide signatures in a single gene dataset (1). However the same is not true of the phylogenetic classification of the Archaea, based on marker-genes and reversible evolution models that rely exclusively on point mutations, specifically substitution mutations, which may not be ideal phylogenetic markers (59), especially for resolving the earliest divergences of life.

The Three-domains of Life hypothesis (25), which was initially based on the interpretation of an unrooted rRNA tree (of life) (1), was put forward largely to emphasize the uniqueness of the Archaea, ascribed to an exclusive lineal descent. Although many lines of evidence, molecular or otherwise, support the uniqueness of the Archaea, phylogenetic analysis of genomic signatures does not support the presumed primitive state of Archaea or Bacteria, and the common belief that Archaea and Bacteria are ancestors of Eukarya (1, 10, 40, 85). Models of evolution of genomic features support a Two-domains (or rather two empires) of Life hypothesis (8), as well as the independent origins and parallel descent of eukaryote and akaryote species (9, 13, 28, 86).

## Data and methods

### Data collection and curation

#### Marker domains datasets

Character matrices of homologous protein-domains, coded as binary-state characters were assembled from genome annotations of SCOP-domains available through the SUPERFAMILY HMM library and genome assignments server; v. 1.75 (http://supfam.org/SUPERFAMILY/) (55, 87).

(i) 141-species dataset was obtained from a previous study (28)

(ii) The 141-species dataset was updated with representatives of novel species described recently, largely with archaeal species from TACK group (29), DPANN group (4) and Asgard group including the Lokiarchaeota (18). In addition, species sampling was enhanced with representatives from the candidate phyla (unclassified) described for bacterial species and with unicellular species of eukaryotes, to a total of 222 species. The complete list of the species with their respective Taxonomy IDs is available in SI Table 1.

When genome annotations were unavailable from SUPERFAMILY database, curated reference proteomes were obtained from the universal protein resource (http://www.uniprot.org/proteomes/). SCOP-domains were annotated using the HMM library and genome annotation tools and routines recommended by the SUPERFAMILY resource.

#### Marker genes datasets

Marker gene datasets from previous studies were obtained as follows, (i) 29 core-genes alignment (16) and (ii) SSU rRNA alignment and 48 core-genes alignments (18).

### Exploratory data analysis

DDNs were constructed with SplitsTree v. 4.14. Split networks were computed using the NeighborNet method from the observed P-distances of the taxa for both nucleotide- and amino acid-characters. Split networks of the protein-domain characterss were computed from Hamming distance, which is identical to the P-distance. The networks were drawn with the equal angle algorithm.

### Phylogenetic analyses

Concatenated gene tree inference: Extensive analyses of the concatenated core-genes datasets are reported in the original studies (16, 18). Analysis here was restricted to the 29 core-genes dataset due its relatively small taxon sampling (44 species) compared to the 48 core-genes dataset (96 species) since there is little difference in data quality, but the computational time/resources required is significantly lesser. Moreover, the general conclusions based on these datasets are consistent despite a smaller taxon sampling, particularly of archaeal species (26 as opposed to 64 in the larger sampling).

Best-fitting amino acid substitution models were chosen using Smart Model Selection (SMS) (88) compatible with PhyML tree inference methods (89). Trees were estimated with a rate-homogeneous LG model as well as rate-heterogeneous versions of the LG model. Site-specific rate variation was approximated using the gamma distribution with 4, 8 and 12 rate categories, LG+G4, LG+G8 and LG+G12, respectively. More complex models (SI Table 2) that account for invariable sites (LG+GX+I) and/or models that compute alignment-specific state frequencies (LG+GX+F) were also used, but the trees inferred were identical to trees estimated from LG+GX models, and therefore not reported here. Log likelihoods ratio (LLR) was calculated as the difference in the raw log likelihoods for each model.

Genome tree inference: The Mk model (35) is the most widely implemented model for phylogenetic inference in the probabilistic framework (maximum likelihood (ML) and Bayesian methods) applicable to complex features coded as binary characters. However, only the reversible model is implemented in ML methods at present. Both reversible and directional evolution models as well as model selection routines implemented in MrBayes 3.2 (43, 90) were used. Two independent runs of Metropolis-Coupled MCMC algorithm was used with four chains each, sampling every 500th generation. The first half of the generations was discarded as burn-in. MCMC sampling was run until convergence, unless mentioned otherwise. Convergence was assessed through the average standard deviation of split frequencies (ASDSF, less than 0.01) for tree topology and the potential scale reduction factor (PSRF, equal to 1.00) for scalar parameters, unless mentioned otherwise. Bayes factors for model comparison were calculated using the harmonic mean estimator in MrBayes. The log Bayes factor (LBF) was calculated as the difference in the log likelihoods for each model.

Convergence between independent runs was generally slower for directional models compared to the reversible models. When convergence was extremely slow (requiring more than 100 million generations and/or more than 15 days run-time) topology constraints corresponding to the clusters derived in the unrooted trees (Fig. 3E) were applied to improve convergence rates. In general these clusters/constraints corresponded to named taxonomic groups e.g. Fungi, Metazoa, Crenarchaeota, etc. Convergence assessment between independent runs was relaxed for three specific cases that did not converge sufficiently at the time of submission: the unrooted tree with Mk-uniform-rates model (ASDSF 0.05; PSRF 1.03), rooted trees corresponding to root-R2 (ASDSF 0.5; PSRF 1.04) and root-R3 (ASDSF 0.029; PSRF 1.03). In the three cases specified, the difference in bipartitions is in the shallow parts (minor branches) of the tree, but not within the major branches. For assessing well-supported major branches of the tree, ASDSF values between 0.01 and 0.05 may be adequate, as recommended by the authors (91).

## Funding

This research received no specific grant from any funding agency in the public, commercial, or not-for-profit sectors. Work by this author was partially supported by The Swedish Research Council (to Måns Ehrenberg) and the Knut and Alice Wallenberg Foundation, RiboCORE (to Måns Ehrenberg and Dan Andersson).

I am grateful to Charles (Chuck) Kurland and Måns Ehrenberg for support and encouragement. I thank Chuck Kurland and Siv Andersson for the discussions in general; Chuck for the many stimulating debates and Siv for inspiring the article title, in part; Seraina Klopfstein for providing the algorithms for implementing the directional model in MrBayes and for helpful suggestions and Erling Wikman for help with computing equipment.

## Supplementary Information

**SI Table 1.**
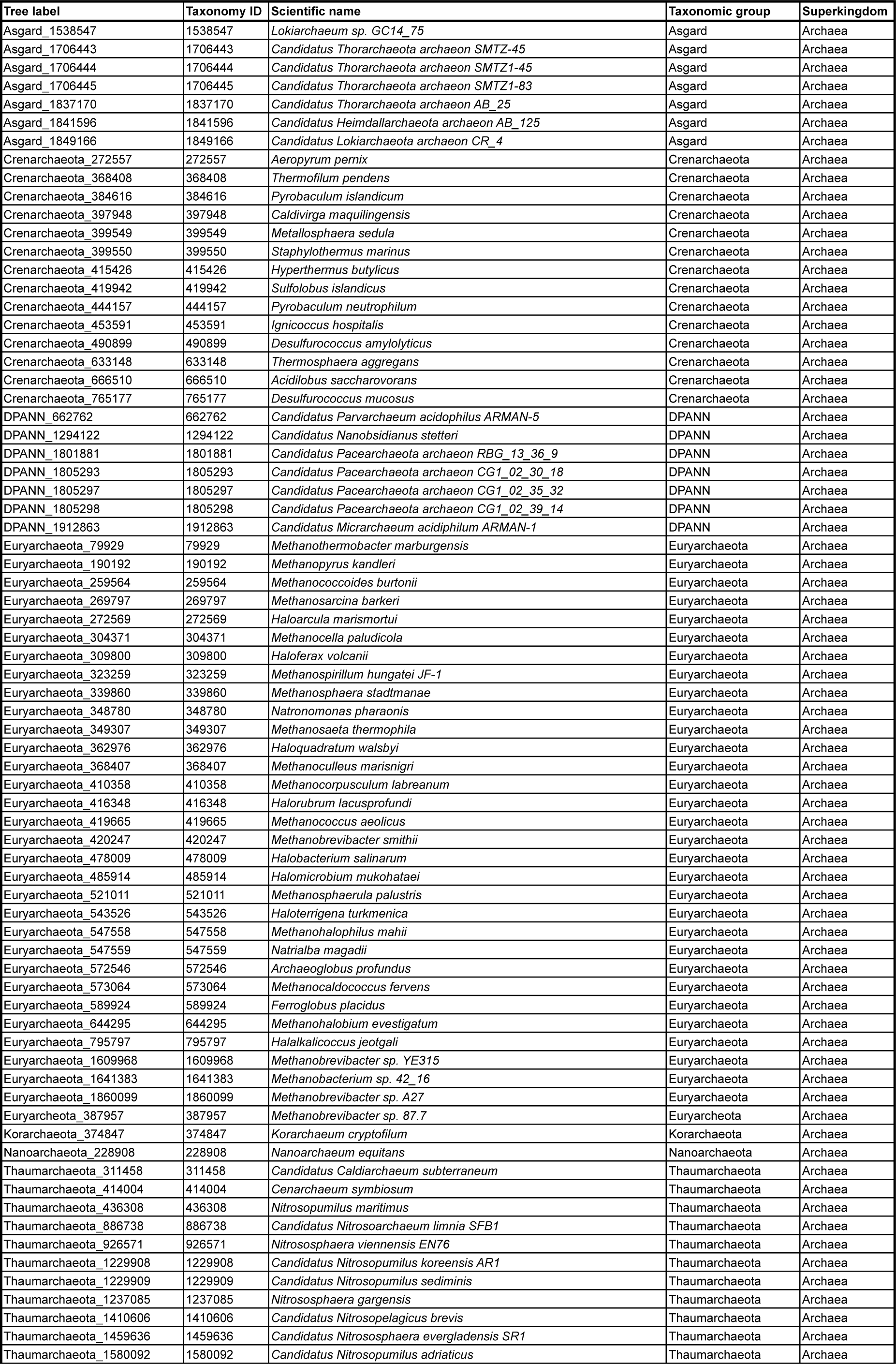
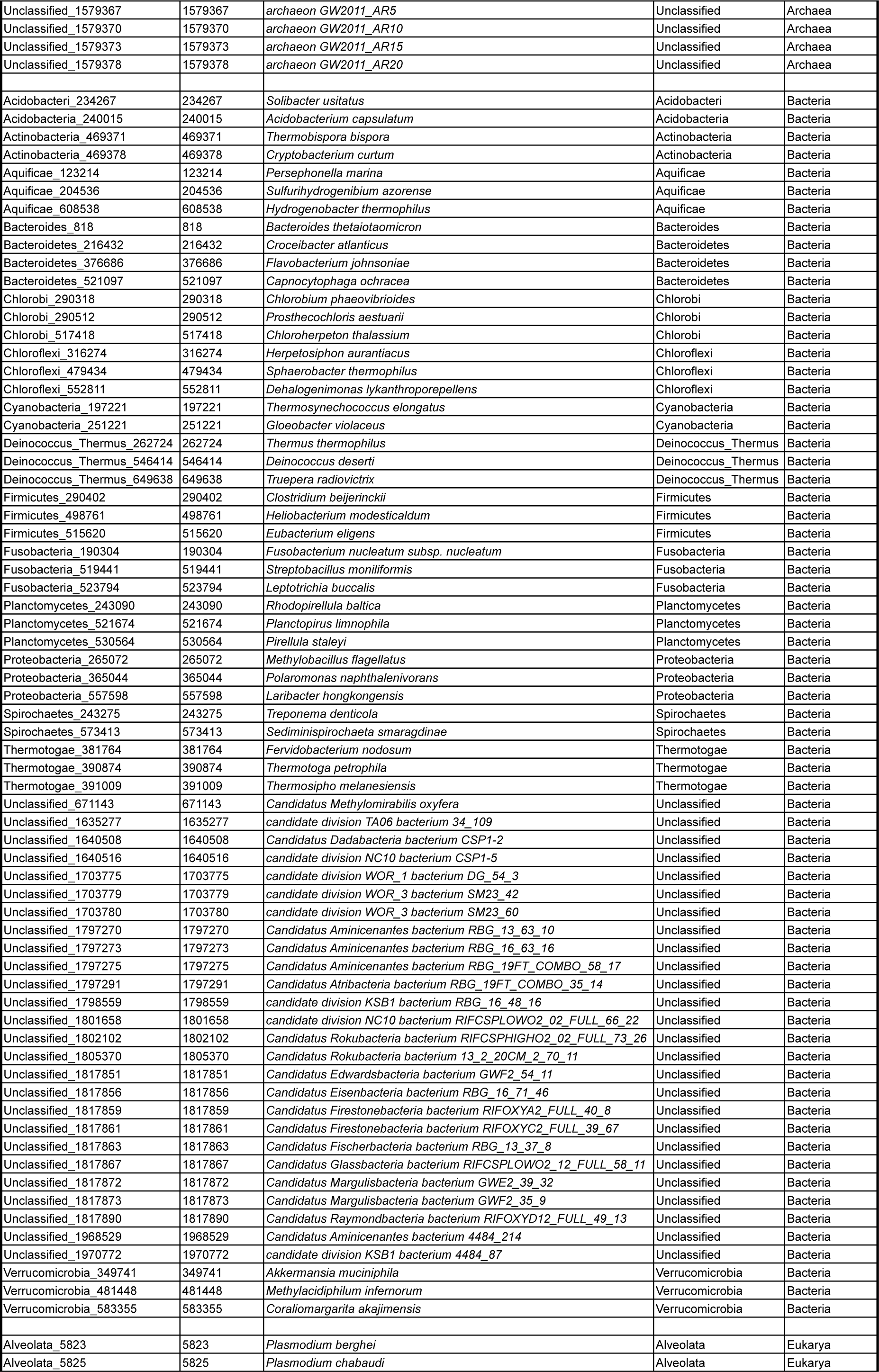
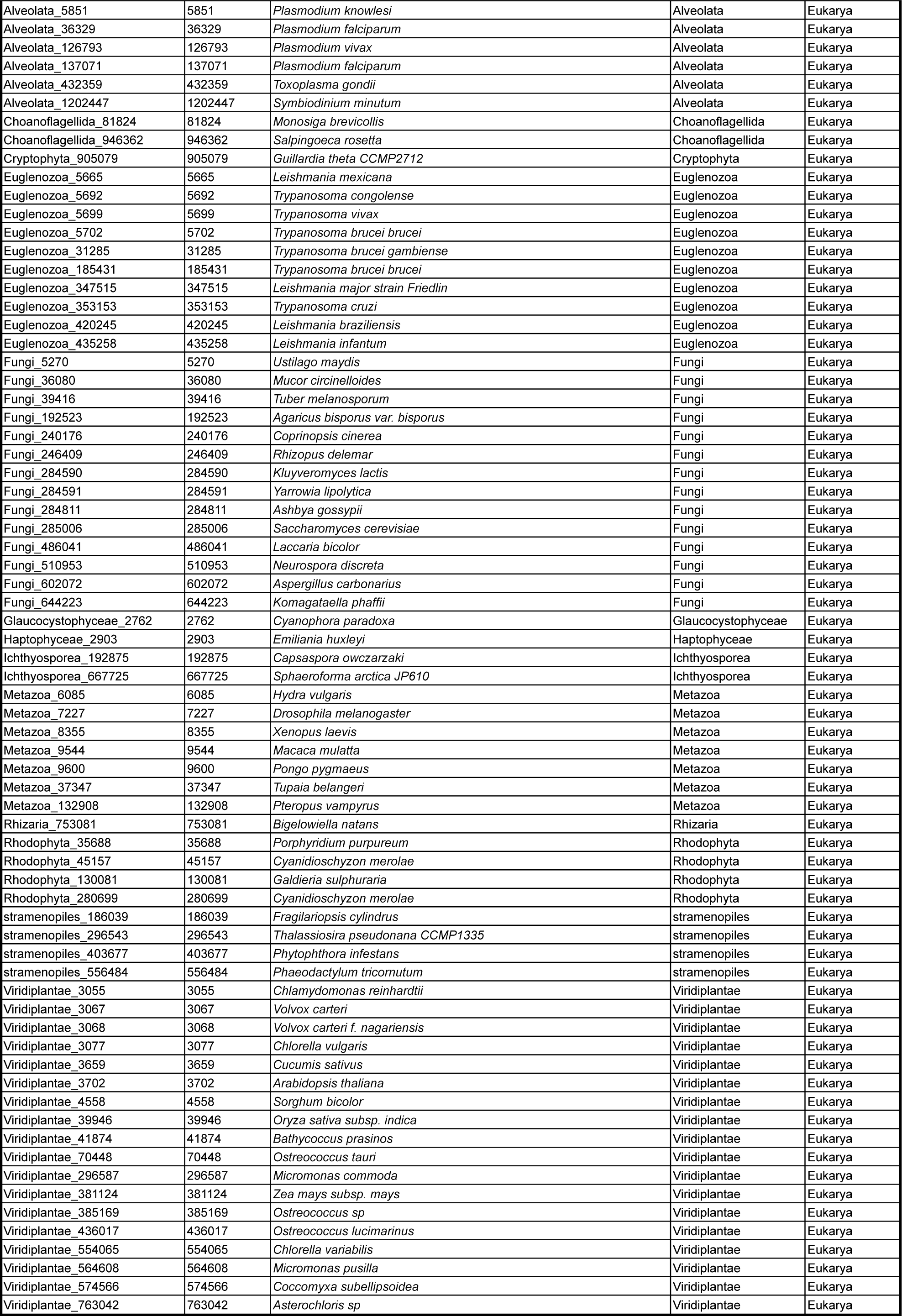
List of organisms analyzed in Fig. 2B, Fig. 3D,E as well as Fig. 5A

| Tree label | Taxonomy ID | Scientific name | Taxonomic group | Superkingdom |
| --- | --- | --- | --- | --- |
| Asgard_1538547 | 1538547 | *Lokiarchaeum* sp. GC14_75 | Asgard | Archaea |
| Asgard_1706443 | 1706443 | *Candidatus Thorarchaeota archaeon SMTZ-45* | Asgard | Archaea |
| Asgard_1706444 | 1706444 | *Candidatus Thorarchaeota archaeon SMTZ1-45* | Asgard | Archaea |
| Asgard_1706445 | 1706445 | *Candidatus Thorarchaeota archaeon SMTZ1-83* | Asgard | Archaea |
| Asgard_1837170 | 1837170 | *Candidatus Thorarchaeota archaeon AB_25* | Asgard | Archaea |
| Asgard_1841596 | 1841596 | *Candidatus Heimdallarchaeota archaeon AB_125* | Asgard | Archaea |
| Asgard_1849166 | 1849166 | *Candidatus Lokiarchaeota archaeon CR_4* | Asgard | Archaea |
| Crenarchaeota_272557 | 272557 | *Aeropyrum pernix* | Crenarchaeota | Archaea |
| Crenarchaeota_368408 | 368408 | *Thermofilum pendens* | Crenarchaeota | Archaea |
| Crenarchaeota_384616 | 384616 | *Pyrobaculum islandicum* | Crenarchaeota | Archaea |
| Crenarchaeota_397948 | 397948 | *Caldivirga maquilingensis* | Crenarchaeota | Archaea |
| Crenarchaeota_399549 | 399549 | *Metallosphaera sedula* | Crenarchaeota | Archaea |
| Crenarchaeota_399550 | 399550 | *Staphylothermus marinus* | Crenarchaeota | Archaea |
| Crenarchaeota_415426 | 415426 | *Hyperthermus butylicus* | Crenarchaeota | Archaea |
| Crenarchaeota_419942 | 419942 | *Sulfolobus islandicus* | Crenarchaeota | Archaea |
| Crenarchaeota_444157 | 444157 | *Pyrobaculum neutrophilum* | Crenarchaeota | Archaea |
| Crenarchaeota_453591 | 453591 | *Ignicoccus hospitalis* | Crenarchaeota | Archaea |
| Crenarchaeota_490899 | 490899 | *Desulfurococcus amylolyticus* | Crenarchaeota | Archaea |
| Crenarchaeota_633148 | 633148 | *Thermosphaera aggregans* | Crenarchaeota | Archaea |
| Crenarchaeota_666510 | 666510 | *Acidilobus saccharovorans* | Crenarchaeota | Archaea |
| Crenarchaeota_765177 | 765177 | *Desulfurococcus mucosus* | Crenarchaeota | Archaea |
| DPANN_662762 | 662762 | *Candidatus Parvarchaeum acidophilum* ARMAN-5 | DPANN | Archaea |
| DPANN_1294122 | 1294122 | *Candidatus Nanobsidianus stetteri* | DPANN | Archaea |
| DPANN_1801881 | 1801881 | *Candidatus Pacearchaeota archaeon RBG_13_36_9* | DPANN | Archaea |
| DPANN_1805293 | 1805293 | *Candidatus Pacearchaeota archaeon CG1_02_30_18* | DPANN | Archaea |
| DPANN_1805297 | 1805297 | *Candidatus Pacearchaeota archaeon CG1_02_35_32* | DPANN | Archaea |
| DPANN_1805298 | 1805298 | *Candidatus Pacearchaeota archaeon CG1_02_39_14* | DPANN | Archaea |
| DPANN_1912863 | 1912863 | *Candidatus Micrarchaeum acidiphilum* ARMAN-1 | DPANN | Archaea |
| Euryarchaeota_79929 | 79929 | *Methanothermobacter marburgensis* | Euryarchaeota | Archaea |
| Euryarchaeota_190192 | 190192 | *Methanopyrus kandleri* | Euryarchaeota | Archaea |
| Euryarchaeota_259564 | 259564 | *Methanococcoides burtonii* | Euryarchaeota | Archaea |
| Euryarchaeota_269797 | 269797 | *Methanosarcina barkeri* | Euryarchaeota | Archaea |
| Euryarchaeota_272569 | 272569 | *Haloarcula marismortui* | Euryarchaeota | Archaea |
| Euryarchaeota_304371 | 304371 | *Methanocella paludicola* | Euryarchaeota | Archaea |
| Euryarchaeota_309800 | 309800 | *Haloferax volcanii* | Euryarchaeota | Archaea |
| Euryarchaeota_323259 | 323259 | *Methanospirillum hungatei* JF-1 | Euryarchaeota | Archaea |
| Euryarchaeota_339860 | 339860 | *Methanosphaera stadtmanae* | Euryarchaeota | Archaea |
| Euryarchaeota_348780 | 348780 | *Natronomonas pharaonis* | Euryarchaeota | Archaea |
| Euryarchaeota_349307 | 349307 | *Methanosaeta thermophila* | Euryarchaeota | Archaea |
| Euryarchaeota_362976 | 362976 | *Haloquadratum walsbyi* | Euryarchaeota | Archaea |
| Euryarchaeota_368407 | 368407 | *Methanoculleus marisnigri* | Euryarchaeota | Archaea |
| Euryarchaeota_410358 | 410358 | *Methanocorpusculum labreanum* | Euryarchaeota | Archaea |
| Euryarchaeota_416348 | 416348 | *Halorubrum lacusprofundi* | Euryarchaeota | Archaea |
| Euryarchaeota_419665 | 419665 | *Methanococcus aeolicus* | Euryarchaeota | Archaea |
| Euryarchaeota_420247 | 420247 | *Methanobrevibacter smithii* | Euryarchaeota | Archaea |
| Euryarchaeota_478009 | 478009 | *Halobacterium salinarum* | Euryarchaeota | Archaea |
| Euryarchaeota_485914 | 485914 | *Halomicrobium mukohataei* | Euryarchaeota | Archaea |
| Euryarchaeota_521011 | 521011 | *Methanosphaerula palustris* | Euryarchaeota | Archaea |
| Euryarchaeota_543526 | 543526 | *Haloterrigena turkmenica* | Euryarchaeota | Archaea |
| Euryarchaeota_547558 | 547558 | *Methanohalophilus mahii* | Euryarchaeota | Archaea |
| Euryarchaeota_547559 | 547559 | *Natrialba magadii* | Euryarchaeota | Archaea |
| Euryarchaeota_572546 | 572546 | *Archaeoglobus profundus* | Euryarchaeota | Archaea |
| Euryarchaeota_573064 | 573064 | *Methanocaldococcus fervens* | Euryarchaeota | Archaea |
| Euryarchaeota_589924 | 589924 | *Ferroglobus placidus* | Euryarchaeota | Archaea |
| Euryarchaeota_644295 | 644295 | *Methanohalobium evestigatum* | Euryarchaeota | Archaea |
| Euryarchaeota_795797 | 795797 | *Halalkalicoccus jeotgali* | Euryarchaeota | Archaea |
| Euryarchaeota_1609968 | 1609968 | *Methanobrevibacter* sp. YE315 | Euryarchaeota | Archaea |
| Euryarchaeota_1641383 | 1641383 | *Methanobacterium* sp. 42_16 | Euryarchaeota | Archaea |
| Euryarchaeota_1860099 | 1860099 | *Methanobrevibacter* sp. A27 | Euryarchaeota | Archaea |
| Euryarchaeota_387957 | 387957 | *Methanobrevibacter* sp. 87.7 | Euryarchaeota | Archaea |
| Korarchaeota_374847 | 374847 | *Korarchaeum cryptofilum* | Korarchaeota | Archaea |
| Nanoarchaeota_228908 | 228908 | *Nanoarchaeum equitans* | Nanoarchaeota | Archaea |
| Thaumarchaeota_311458 | 311458 | *Candidatus Caldiarchaeum subterraneum* | Thaumarchaeota | Archaea |
| Thaumarchaeota_414004 | 414004 | *Cenarchaeum symbiosum* | Thaumarchaeota | Archaea |
| Thaumarchaeota_436308 | 436308 | *Nitrosopumilus maritimus* | Thaumarchaeota | Archaea |
| Thaumarchaeota_886738 | 886738 | *Candidatus Nitrosoarchaeum limnia* SFB1 | Thaumarchaeota | Archaea |
| Thaumarchaeota_926571 | 926571 | *Nitrososphaera viennensis* EN76 | Thaumarchaeota | Archaea |
| Thaumarchaeota_1229908 | 1229908 | *Candidatus Nitrosopumilus koreensis* AR1 | Thaumarchaeota | Archaea |
| Thaumarchaeota_1229909 | 1229909 | *Candidatus Nitrosopumilus sediminis* | Thaumarchaeota | Archaea |
| Thaumarchaeota_1237085 | 1237085 | *Nitrososphaera gargensis* | Thaumarchaeota | Archaea |
| Thaumarchaeota_1410606 | 1410606 | *Candidatus Nitrosopelagicus brevis* | Thaumarchaeota | Archaea |
| Thaumarchaeota_1459636 | 1459636 | *Candidatus Nitrososphaera evergladensis* SR1 | Thaumarchaeota | Archaea |
| Thaumarchaeota_1580092 | 1580092 | *Candidatus Nitrosopumilus adriaticus* | Thaumarchaeota | Archaea |
| Unclassified_1579367 | 1579367 | archaeon GW2011_AR5 | Unclassified | Archaea |
| Unclassified_1579370 | 1579370 | archaeon GW2011_AR10 | Unclassified | Archaea |
| Unclassified_1579373 | 1579373 | archaeon GW2011_AR15 | Unclassified | Archaea |
| Unclassified_1579378 | 1579378 | archaeon GW2011_AR20 | Unclassified | Archaea |
| Acidobacteri_234267 | 234267 | *Solibacter usitatus* | Acidobacteri | Bacteria |
| Acidobacteria_240015 | 240015 | *Acidobacterium capsulatum* | Acidobacteria | Bacteria |
| Actinobacteria_469371 | 469371 | *Thermobispora bispora* | Actinobacteria | Bacteria |
| Actinobacteria_469378 | 469378 | *Cryptobacterium curtum* | Actinobacteria | Bacteria |
| Aquificae_123214 | 123214 | *Persephonella marina* | Aquificae | Bacteria |
| Aquificae_204536 | 204536 | *Sulfurihydrogenibium azorense* | Aquificae | Bacteria |
| Aquificae_608538 | 608538 | *Hydrogenobacter thermophilus* | Aquificae | Bacteria |
| Bacteroides_818 | 818 | *Bacteroides thetaiotaomicron* | Bacteroides | Bacteria |
| Bacteroidetes_216432 | 216432 | *Croceibacter atlanticus* | Bacteroidetes | Bacteria |
| Bacteroidetes_376686 | 376686 | *Flavobacterium johnsoniae* | Bacteroidetes | Bacteria |
| Bacteroidetes_521097 | 521097 | *Capnocytophaga ochracea* | Bacteroidetes | Bacteria |
| Chlorobi_290318 | 290318 | *Chlorobium phaeovibrioides* | Chlorobi | Bacteria |
| Chlorobi_290512 | 290512 | *Prosthecochloris aestuarii* | Chlorobi | Bacteria |
| Chlorobi_517418 | 517418 | *Chloroherpeton thalassium* | Chlorobi | Bacteria |
| Chloroflexi_316274 | 316274 | *Herpetosiphon aurantiacus* | Chloroflexi | Bacteria |
| Chloroflexi_479434 | 479434 | *Sphaerobacter thermophilus* | Chloroflexi | Bacteria |
| Chloroflexi_552811 | 552811 | *Dehalogenimonas lykanthroporepellens* | Chloroflexi | Bacteria |
| Cyanobacteria_197221 | 197221 | *Thermosynechococcus elongatus* | Cyanobacteria | Bacteria |
| Cyanobacteria_251221 | 251221 | *Gloeobacter violaceus* | Cyanobacteria | Bacteria |
| Deinococcus_Thermus_262724 | 262724 | *Thermus thermophilus* | Deinococcus_Thermus | Bacteria |
| Deinococcus_Thermus_546414 | 546414 | *Deinococcus deserti* | Deinococcus_Thermus | Bacteria |
| Deinococcus_Thermus_649638 | 649638 | *Truepera radiovictrix* | Deinococcus_Thermus | Bacteria |
| Firmicutes_290402 | 290402 | *Clostridium beijerinckii* | Firmicutes | Bacteria |
| Firmicutes_498761 | 498761 | *Helibacterium modesticaldum* | Firmicutes | Bacteria |
| Firmicutes_515620 | 515620 | *Eubacterium eligens* | Firmicutes | Bacteria |
| Fusobacteria_190304 | 190304 | *Fusobacterium nucleatum* subsp. *nucleatum* | Fusobacteria | Bacteria |
| Fusobacteria_519441 | 519441 | *Streptobacillus moniliformis* | Fusobacteria | Bacteria |
| Fusobacteria_523794 | 523794 | *Leptotrichia buccalis* | Fusobacteria | Bacteria |
| Planctomycetes_243090 | 243090 | *Rhodopirellula baltica* | Planctomycetes | Bacteria |
| Planctomycetes_521674 | 521674 | *Planctopirax limnophila* | Planctomycetes | Bacteria |
| Planctomycetes_530564 | 530564 | *Pirellula staleyi* | Planctomycetes | Bacteria |
| Proteobacteria_265072 | 265072 | *Methylobacillus flagellatus* | Proteobacteria | Bacteria |
| Proteobacteria_365044 | 365044 | *Polaromonas naphthalenivorans* | Proteobacteria | Bacteria |
| Proteobacteria_557598 | 557598 | *Laribacter hongkongensis* | Proteobacteria | Bacteria |
| Spirochaetes_243275 | 243275 | *Treponema denticola* | Spirochaetes | Bacteria |
| Spirochaetes_573413 | 573413 | *Sediminispirochaeta smaragdinae* | Spirochaetes | Bacteria |
| Thermotogae_381764 | 381764 | *Fervidobacterium nodosum* | Thermotogae | Bacteria |
| Thermotogae_390874 | 390874 | *Thermotoga petrophila* | Thermotogae | Bacteria |
| Thermotogae_391009 | 391009 | *Thermosiphon melanesiensis* | Thermotogae | Bacteria |
| Unclassified_671143 | 671143 | *Candidatus Methyloirabialis oxyfera* | Unclassified | Bacteria |
| Unclassified_1635277 | 1635277 | candidate division TA06 bacterium 34_109 | Unclassified | Bacteria |
| Unclassified_1640508 | 1640508 | *Candidatus Dadabacteria bacterium CSP1-2* | Unclassified | Bacteria |
| Unclassified_1640516 | 1640516 | candidate division NC10 bacterium CSP1-5 | Unclassified | Bacteria |
| Unclassified_1703775 | 1703775 | candidate division WOR_1 bacterium DG_54_3 | Unclassified | Bacteria |
| Unclassified_1703779 | 1703779 | candidate division WOR_3 bacterium SM23_42 | Unclassified | Bacteria |
| Unclassified_1703780 | 1703780 | candidate division WOR_3 bacterium SM23_60 | Unclassified | Bacteria |
| Unclassified_1797270 | 1797270 | *Candidatus Aminicenantes bacterium RBG_13_63_10* | Unclassified | Bacteria |
| Unclassified_1797273 | 1797273 | *Candidatus Aminicenantes bacterium RBG_16_63_16* | Unclassified | Bacteria |
| Unclassified_1797275 | 1797275 | *Candidatus Aminicenantes bacterium RBG_19FT_COMBO_58_17* | Unclassified | Bacteria |
| Unclassified_1797291 | 1797291 | *Candidatus Atribacteria bacterium RBG_19FT_COMBO_35_14* | Unclassified | Bacteria |
| Unclassified_1798559 | 1798559 | candidate division KSB1 bacterium RBG_16_48_16 | Unclassified | Bacteria |
| Unclassified_1801658 | 1801658 | candidate division NC10 bacterium RIFCSPLOWO2_02_FULL_66_22 | Unclassified | Bacteria |
| Unclassified_1802102 | 1802102 | *Candidatus Rokubacteria bacterium RIFCSPHIGHO2_02_FULL_73_26* | Unclassified | Bacteria |
| Unclassified_1805370 | 1805370 | *Candidatus Rokubacteria bacterium 13_2_20CM_2_70_11* | Unclassified | Bacteria |
| Unclassified_1817851 | 1817851 | *Candidatus Edwardsbacteria bacterium GWF2_54_11* | Unclassified | Bacteria |
| Unclassified_1817856 | 1817856 | *Candidatus Eisenbacteria bacterium RBG_16_71_46* | Unclassified | Bacteria |
| Unclassified_1817859 | 1817859 | *Candidatus Firestonebacteria bacterium RIFOXYA2_FULL_40_8* | Unclassified | Bacteria |
| Unclassified_1817861 | 1817861 | *Candidatus Firestonebacteria bacterium RIFOXYC2_FULL_39_67* | Unclassified | Bacteria |
| Unclassified_1817863 | 1817863 | *Candidatus Fischerbacteria bacterium RBG_13_37_8* | Unclassified | Bacteria |
| Unclassified_1817867 | 1817867 | *Candidatus Glassbacteria bacterium RIFCSPLOWO2_12_FULL_58_11* | Unclassified | Bacteria |
| Unclassified_1817872 | 1817872 | *Candidatus Margulisbacteria bacterium GWE2_39_32* | Unclassified | Bacteria |
| Unclassified_1817873 | 1817873 | *Candidatus Margulisbacteria bacterium GWF2_35_9* | Unclassified | Bacteria |
| Unclassified_1817890 | 1817890 | *Candidatus Raymondobacteria bacterium RIFOXYD12_FULL_49_13* | Unclassified | Bacteria |
| Unclassified_1968529 | 1968529 | *Candidatus Aminicenantes bacterium 4484_214* | Unclassified | Bacteria |
| Unclassified_1970772 | 1970772 | candidate division KSB1 bacterium 4484_87 | Unclassified | Bacteria |
| Verrucomicrobia_349741 | 349741 | *Akkermansia muciniphila* | Verrucomicrobia | Bacteria |
| Verrucomicrobia_481448 | 481448 | *Methylacidiphilum inferorum* | Verrucomicrobia | Bacteria |
| Verrucomicrobia_583355 | 583355 | *Coralimargarita akajimensis* | Verrucomicrobia | Bacteria |
| Alveolata_5823 | 5823 | *Plasmodium berghei* | Alveolata | Eukarya |
| Alveolata_5825 | 5825 | *Plasmodium chabaudi* | Alveolata | Eukarya |
| Alveolata_5851 | 5851 | *Plasmodium knowlesi* | Alveolata | Eukarya |
| Alveolata_36329 | 36329 | *Plasmodium falciparum* | Alveolata | Eukarya |
| Alveolata_126793 | 126793 | *Plasmodium vivax* | Alveolata | Eukarya |
| Alveolata_137071 | 137071 | *Plasmodium falciparum* | Alveolata | Eukarya |
| Alveolata_432359 | 432359 | *Toxoplasma gondii* | Alveolata | Eukarya |
| Alveolata_1202447 | 1202447 | *Symbiodinium minutum* | Alveolata | Eukarya |
| Choanoflagellida_81824 | 81824 | *Monosiga brevicollis* | Choanoflagellida | Eukarya |
| Choanoflagellida_946362 | 946362 | *Salpingoeca rosetta* | Choanoflagellida | Eukarya |
| Cryptophyta_905079 | 905079 | *Guillardia theta CCMP2712* | Cryptophyta | Eukarya |
| Euglenozoa_5665 | 5665 | *Leishmania mexicana* | Euglenozoa | Eukarya |
| Euglenozoa_5692 | 5692 | *Trypanosoma congolense* | Euglenozoa | Eukarya |
| Euglenozoa_5699 | 5699 | *Trypanosoma vivax* | Euglenozoa | Eukarya |
| Euglenozoa_5702 | 5702 | *Trypanosoma brucei brucei* | Euglenozoa | Eukarya |
| Euglenozoa_31285 | 31285 | *Trypanosoma brucei gambiense* | Euglenozoa | Eukarya |
| Euglenozoa_185431 | 185431 | *Trypanosoma brucei brucei* | Euglenozoa | Eukarya |
| Euglenozoa_347515 | 347515 | *Leishmania major strain Friedlin* | Euglenozoa | Eukarya |
| Euglenozoa_353153 | 353153 | *Trypanosoma cruzi* | Euglenozoa | Eukarya |
| Euglenozoa_420245 | 420245 | *Leishmania braziliensis* | Euglenozoa | Eukarya |
| Euglenozoa_435258 | 435258 | *Leishmania infantum* | Euglenozoa | Eukarya |
| Fungi_5270 | 5270 | *Ustilago maydis* | Fungi | Eukarya |
| Fungi_36080 | 36080 | *Mucor circinelloides* | Fungi | Eukarya |
| Fungi_39416 | 39416 | *Tuber melanosporum* | Fungi | Eukarya |
| Fungi_192523 | 192523 | *Agaricus bisporus var. bisporus* | Fungi | Eukarya |
| Fungi_240176 | 240176 | *Coprinopsis cinerea* | Fungi | Eukarya |
| Fungi_246409 | 246409 | *Rhizopus delemar* | Fungi | Eukarya |
| Fungi_284590 | 284590 | *Kluyveromyces lactis* | Fungi | Eukarya |
| Fungi_284591 | 284591 | *Yarrowia lipolytica* | Fungi | Eukarya |
| Fungi_284811 | 284811 | *Ashbya gossypii* | Fungi | Eukarya |
| Fungi_285006 | 285006 | *Saccharomyces cerevisiae* | Fungi | Eukarya |
| Fungi_486041 | 486041 | *Laccaria bicolor* | Fungi | Eukarya |
| Fungi_510953 | 510953 | *Neurospora discreta* | Fungi | Eukarya |
| Fungi_602072 | 602072 | *Aspergillus carbonarius* | Fungi | Eukarya |
| Fungi_644223 | 644223 | *Komagataella phaffii* | Fungi | Eukarya |
| Glaucozystophyceae_2762 | 2762 | *Cyanophora paradoxa* | Glaucozystophyceae | Eukarya |
| Haptophyceae_2903 | 2903 | *Emiliania huxleyi* | Haptophyceae | Eukarya |
| Ichthyosporea_192875 | 192875 | *Capsaspora owczarzaki* | Ichthyosporea | Eukarya |
| Ichthyosporea_667725 | 667725 | *Sphaeroforma arctica JP610* | Ichthyosporea | Eukarya |
| Metazoa_6085 | 6085 | *Hydra vulgaris* | Metazoa | Eukarya |
| Metazoa_7227 | 7227 | *Drosophila melanogaster* | Metazoa | Eukarya |
| Metazoa_8355 | 8355 | *Xenopus laevis* | Metazoa | Eukarya |
| Metazoa_9544 | 9544 | *Macaca mulatta* | Metazoa | Eukarya |
| Metazoa_9600 | 9600 | *Pongo pygmaeus* | Metazoa | Eukarya |
| Metazoa_37347 | 37347 | *Tupaia belangeri* | Metazoa | Eukarya |
| Metazoa_132908 | 132908 | *Pteropus vampyrus* | Metazoa | Eukarya |
| Rhizaria_753081 | 753081 | *Bigeloviella natans* | Rhizaria | Eukarya |
| Rhodophyta_35688 | 35688 | *Porphyridium purpureum* | Rhodophyta | Eukarya |
| Rhodophyta_45157 | 45157 | *Cyanidioschyzon merolae* | Rhodophyta | Eukarya |
| Rhodophyta_130081 | 130081 | *Galdieria sulphuraria* | Rhodophyta | Eukarya |
| Rhodophyta_280699 | 280699 | *Cyanidioschyzon merolae* | Rhodophyta | Eukarya |
| stramenopiles_186039 | 186039 | *Fragilariaopsis cylindrus* | stramenopiles | Eukarya |
| stramenopiles_296543 | 296543 | *Thalassiosira pseudonana CCMP1335* | stramenopiles | Eukarya |
| stramenopiles_403677 | 403677 | *Phytophthora infestans* | stramenopiles | Eukarya |
| stramenopiles_556484 | 556484 | *Phaeodactylum tricornutum* | stramenopiles | Eukarya |
| Viridiplantae_3055 | 3055 | *Chlamydomonas reinhardtii* | Viridiplantae | Eukarya |
| Viridiplantae_3067 | 3067 | *Volvox carteri* | Viridiplantae | Eukarya |
| Viridiplantae_3068 | 3068 | *Volvox carteri f. nagariensis* | Viridiplantae | Eukarya |
| Viridiplantae_3077 | 3077 | *Chlorella vulgaris* | Viridiplantae | Eukarya |
| Viridiplantae_3659 | 3659 | *Cucumis sativus* | Viridiplantae | Eukarya |
| Viridiplantae_3702 | 3702 | *Arabidopsis thaliana* | Viridiplantae | Eukarya |
| Viridiplantae_4558 | 4558 | *Sorghum bicolor* | Viridiplantae | Eukarya |
| Viridiplantae_39946 | 39946 | *Oryza sativa subsp. indica* | Viridiplantae | Eukarya |
| Viridiplantae_41874 | 41874 | *Bathycoccus prasinos* | Viridiplantae | Eukarya |
| Viridiplantae_70448 | 70448 | *Ostreococcus tauri* | Viridiplantae | Eukarya |
| Viridiplantae_296587 | 296587 | *Micromonas commoda* | Viridiplantae | Eukarya |
| Viridiplantae_381124 | 381124 | *Zea mays subsp. mays* | Viridiplantae | Eukarya |
| Viridiplantae_385169 | 385169 | *Ostreococcus sp* | Viridiplantae | Eukarya |
| Viridiplantae_436017 | 436017 | *Ostreococcus lucimarinus* | Viridiplantae | Eukarya |
| Viridiplantae_554065 | 554065 | *Chlorella variabilis* | Viridiplantae | Eukarya |
| Viridiplantae_564608 | 564608 | *Micromonas pusilla* | Viridiplantae | Eukarya |
| Viridiplantae_574566 | 574566 | *Coccomyxa subellipsoidea* | Viridiplantae | Eukarya |
| Viridiplantae_763042 | 763042 | *Asterochloris sp* | Viridiplantae | Eukarya |

**SI Table 2.** List of sequence evolution models evaluated for 29 core-genes dataset

| Base Model | Heterogeneity | K | Likelihood | LLR | AIC | BIC |
| --- | --- | --- | --- | --- | --- | --- |
| LG | +G+I | 87 | -492883 | 0 | 985940 | 986553 |
| LG | +G+I+F | 106 | -493145 | 262 | 986502 | 987250 |
| LG | +G | 86 | -493206 | 324 | 986585 | 987191 |
| LG | +G+F | 105 | -493420 | 538 | 987051 | 987791 |
| RtREV | +G+I+F | 106 | -494212 | 1330 | 988637 | 989385 |
| WAG | +G+I+F | 106 | -496011 | 3128 | 992234 | 992982 |
| WAG | +G+I | 87 | -496884 | 4001 | 993942 | 994555 |
| VT | +G+I+F | 106 | -496978 | 4096 | 994169 | 994916 |
| Blosum62 | +G+I | 87 | -497004 | 4121 | 994182 | 994796 |
| VT | +G+I | 87 | -497243 | 4360 | 994660 | 995274 |
| RtREV | +G+I | 87 | -497421 | 4538 | 995016 | 995630 |
| Blosum62 | +G+I+F | 106 | -497536 | 4654 | 995285 | 996033 |
| CpREV | +G+I | 87 | -498666 | 5783 | 997506 | 998120 |
| CpREV | +G+I+F | 106 | -498692 | 5809 | 997596 | 998344 |
| JTT | +G+I | 87 | -501937 | 9055 | 1004049 | 1004662 |
| JTT | +G+I+F | 106 | -501986 | 9103 | 1004183 | 1004931 |
| MtZoa | +G+I+F | 106 | -502729 | 9846 | 1005670 | 1006418 |
| DCMut | +G+I+F | 106 | -503422 | 10539 | 1007055 | 1007803 |
| Dayhoff | +G+I+F | 106 | -503425 | 10543 | 1007063 | 1007811 |
| MtREV | +G+I+F | 106 | -505701 | 12818 | 1011614 | 1012362 |
| MtArt | +G+I+F | 106 | -507424 | 14541 | 1015060 | 1015807 |
| Flu | +G+I+F | 106 | -510433 | 17550 | 1021078 | 1021826 |
| HIVb | +G+I+F | 106 | -514167 | 21284 | 1028546 | 1029294 |
| HIVb | +G+I | 87 | -514445 | 21563 | 1029065 | 1029679 |
| AB | +G+I+F | 106 | -515188 | 22305 | 1030588 | 1031336 |
| MtMam | +G+I+F | 106 | -520429 | 27546 | 1041069 | 1041817 |
| HIVw | +G+I+F | 106 | -532211 | 39328 | 1064634 | 1065381 |

Base Model = Generalized empirical amino acid exchange rate (probability)
Heterogeneity = Parameter for approximating site-specific variation
+GX; where G is discrete Gamma model, X is no. of categories (default X=4)
+I; proportion of invariant sites
+F; empirical amino acid frequencies estimated from the alignment
K = Number of parameters
Likelihood = Raw likelihood score
LLR = Log likelihood ratio given as the difference from the best fitting model
AIC = Akaike information criterion

**SI Fig. 1.**
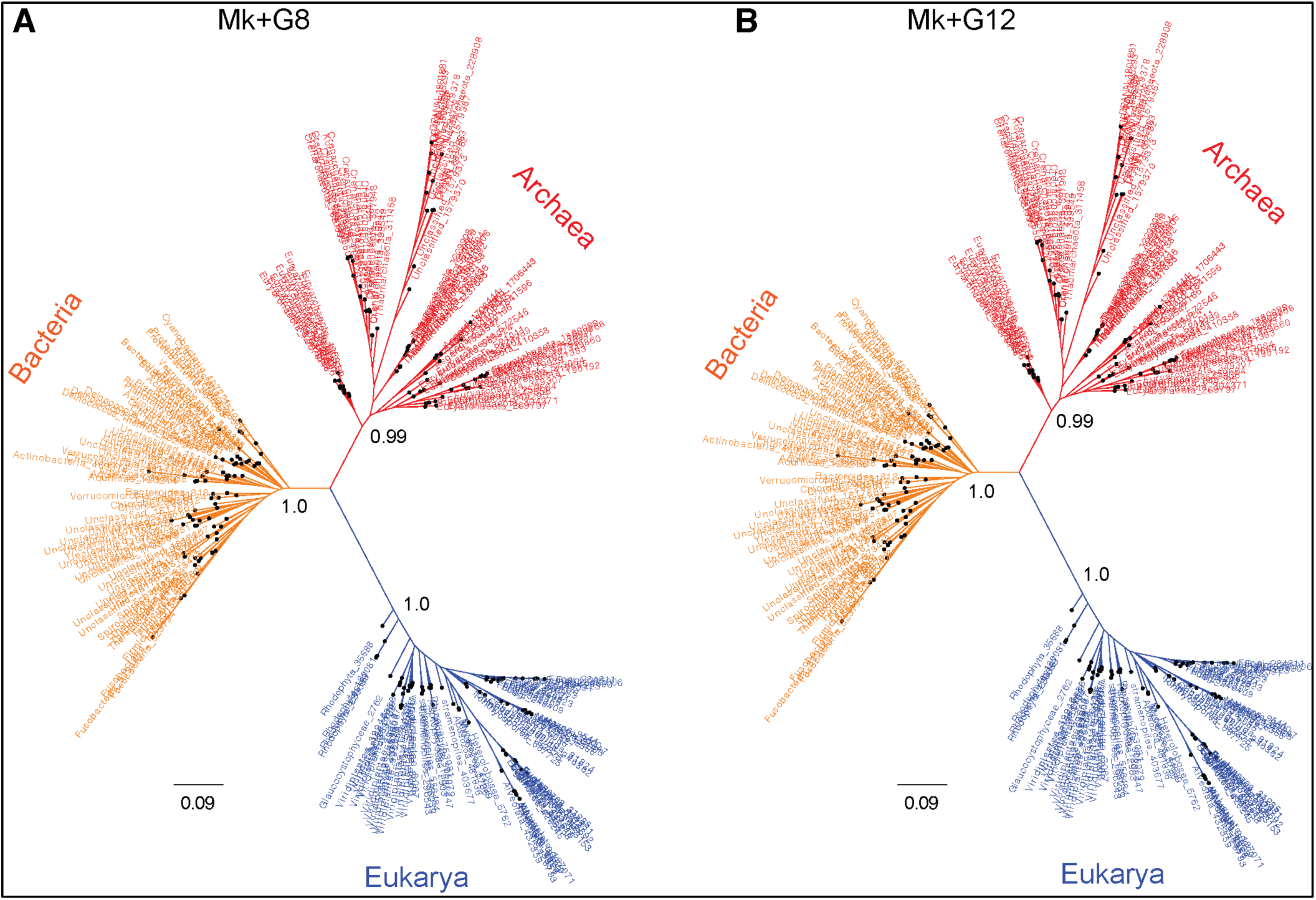
Unrooted genome trees derived from rate-heterogeneous versions of the Mk model. (A) Unrooted tree estimated from Mk+G8 model and (B) from Mk+G12 model. Scale bars represent expected number of changes per character. Branch support (posterior probability) is shown only for the major branches.

